# Identification and mitigation of pervasive off-target activity in CRISPR-Cas9 screens for essential non-coding elements

**DOI:** 10.1101/520569

**Authors:** Josh Tycko, Michael Wainberg, Georgi K. Marinov, Oana Ursu, Gaelen T. Hess, Braeden K. Ego, Aradhana, Amy Li, Alisa Truong, Alexandro E. Trevino, Kaitlyn Spees, David Yao, Irene M. Kaplow, Peyton G. Greenside, David W. Morgens, Douglas H. Phanstiel, Michael P. Snyder, Lacramioara Bintu, William J. Greenleaf, Anshul Kundaje, Michael C. Bassik

## Abstract

Pooled CRISPR-Cas9 screens have recently emerged as a powerful method for functionally characterizing regulatory elements in the non-coding genome, but off-target effects in these experiments have not been systematically evaluated. Here, we conducted a genome-scale screen for essential CTCF loop anchors in the K562 leukemia cell line. Surprisingly, the primary drivers of signal in this screen were single guide RNAs (sgRNAs) with low specificity scores. After removing these guides, we found that there were no CTCF loop anchors critical for cell growth. We also observed this effect in an independent screen fine-mapping the core motifs in enhancers of the *GATA1* gene. We then conducted screens in parallel with CRISPRi and CRISPRa, which do not induce DNA damage, and found that an unexpected and distinct set of off-targets also caused strong confounding growth effects with these epigenome-editing platforms. Promisingly, strict filtering of CRISPRi libraries using GuideScan specificity scores removed these confounded sgRNAs and allowed for the identification of essential enhancers, which we validated extensively. Together, our results show off-target activity can severely limit identification of essential functional motifs by active Cas9, while strictly filtered CRISPRi screens can be reliably used for assaying larger regulatory elements.

## Introduction

Pooled CRISPR-Cas9 screens ^1–5^ have recently emerged as a powerful tool for characterizing the functional importance of genes and non-coding genomic elements. In particular, growth screens have been successfully employed to discover essential genes that determine cell fitness under normal culture conditions ^1,2,6–8^. In addition, CRISPR-Cas9 screens have increasingly been used to functionally characterize the non-coding genome ^9–18^. A variety of approaches have been devised for interrogating non-coding genomic elements. In some instances, active Cas9 nuclease is used to edit candidate functional elements (e.g. transcription factor motifs) at the sequence level by generating indels ^10,19^. Alternatively, the epigenetic environment around a locus can be perturbed using nuclease-dead dCas9 fused to effector domains that can recruit chromatin silencers that modify histones with repressive marks (CRISPRi) ^9,14,20–23^ or activators that recruit transcriptional machinery (CRISPRa) ^11,15,23,24^.

A challenge in interpreting these screens is that CRISPR-Cas9 can bind or edit at unintended off-target genomic sites in a manner that depends on the specificity of the sgRNA sequence ^25–29^. For active Cas9, off-target activity at perfectly matched sites ^30–33^ or sites with 1-2 mismatches ^34,35^ has been shown to reduce cell fitness and confound gene-targeting growth screens. This reduction in cell fitness could be due to accumulating DNA damage from off-target cleavage events. Conversely, for CRISPRi and CRISPRa, the impact of off-target activity on gene-targeting growth screens was shown to be minimal ^3^. However, the impact of off-target activity on screens for essential non-coding regulatory elements has not been studied for any of the three perturbations (active Cas9, CRISPRi and CRISPRa).

To mitigate the impact of off-target effects on screens, sgRNA selection is critical. For gene screens, a large targetable window is present within which all sgRNAs that induce frameshifting indels would be expected to have the same effect on the gene (i.e. a complete knockout), making the selection of highly specific sgRNAs relatively straightforward ^34,36–38^. On the other hand, screens of non-coding elements that use active Cas9 often require the use of lower specificity sgRNAs because regulatory elements, such as individual TF motifs, present a more narrow targeting window from which fewer sgRNAs may be selected.

Despite these challenges, CRISPR-Cas9 screens present an opportunity to systematically perturb and functionally characterize non-coding elements that could not be studied with earlier high-throughput technologies like shRNAs, gene traps, or ORF libraries. One class of candidate *cis*-regulatory elements (ccREs) that have not been functionally dissected in a high-throughput manner are CTCF binding sites in chromatin loop anchors. CTCF binding sites are enriched at the boundaries that partition interphase vertebrate genomes into TADs (Topologically Associated Domains) ^39,40^, and pairs of convergently oriented CTCF motifs are enriched at the anchors of chromatin loops ^40–42^. These chromatin loops and TADs are thought to constrain enhancer-promoter interactions, adding a layer of specificity to the *cis*-regulatory wiring that connects genes with distal regulatory elements. CRISPR-mediated deletions and inversions of individual CTCF sites have been shown to result in reorganization of TADs ^41^ and occasionally in changes in gene expression ^43–45^. Moreover, disruptions of CTCF occupancy have been suggested to be involved in tumorigenesis by leading to pathogenic rewiring of enhancer-promoter interactions ^46–49^. In fact, global degradation of CTCF protein in the cell showed that CTCF is required for the formation and maintenance of TADs and resulted in 370 differentially expressed genes after one day of CTCF depletion ^50^, albeit with only small fold-changes in expression for those genes. However, these type of global perturbations do not reveal the functional importance of individual CTCF sites.

To address this, we set out to perform a genome-wide non-coding screen for essential CTCF binding sites in chromatin loop anchors in the K562 leukemia cell line. We were surprised to discover that the dominant source of signal in our screen was not from deregulated expression of essential genes but was instead consistent with CRISPR-Cas9 off-target activity causing large reductions in cell fitness. This discovery led us to systematically explore the impact of off-target activity across a number of different non-coding screen paradigms. We learned that off-target activity also confounds Cas9 screens for essential functional motifs within enhancers and that CRISPRi/a platforms are similarly vulnerable to off-target activity that significantly reduces cellular fitness. We investigated which non-coding elements can be reliably screened with high-specificity sgRNAs and found that Cas9 screens for essential functional motifs are severely limited by low availability of high-specificity sgRNAs (as determined by a computational specificity score), whereas CRISPRi/a libraries can be properly filtered to avoid confounding off-target activity because their sgRNAs can be selected from a larger targeting window. Together, our results provide principles for the design and interpretation of high-throughput measurements of regulatory element essentiality.

## Results

### CRISPR-Cas9 screens for essential CTCF loop anchors in K562

To identify essential CTCF sites, we performed a Cas9 growth screen with an sgRNA library targeting 4,022 CTCF motifs known to be at loop anchor sites in the K562 cell line according to available Hi-C and CTCF ChIP-seq evidence ^40,51^ (Figure 1A, **Supplementary Table 1**). The library included 2 to 5 sgRNAs per CTCF site that had an expected cleavage site within the motif. The growth effects, measured as guide enrichment from the original sgRNA library plasmid pool to the end of the screen, were highly reproducible between the two independently transduced biological replicates (r^2^ = 0.75, Figure 1B). We observed strong growth effects from the internal positive control sgRNAs that target the exons of essential genes, as well as from sgRNAs targeting the BCR-ABL copy number amplification, which are expected to cause substantial toxicity due to the creation of multiple DNA double-stranded breaks ^30–33,52^. We validated 15 individual sgRNAs using a competitive growth assay, which confirmed the growth effects observed in the pooled screen (r^2^ = 0.69, Figure 1C).

**Figure 1.**
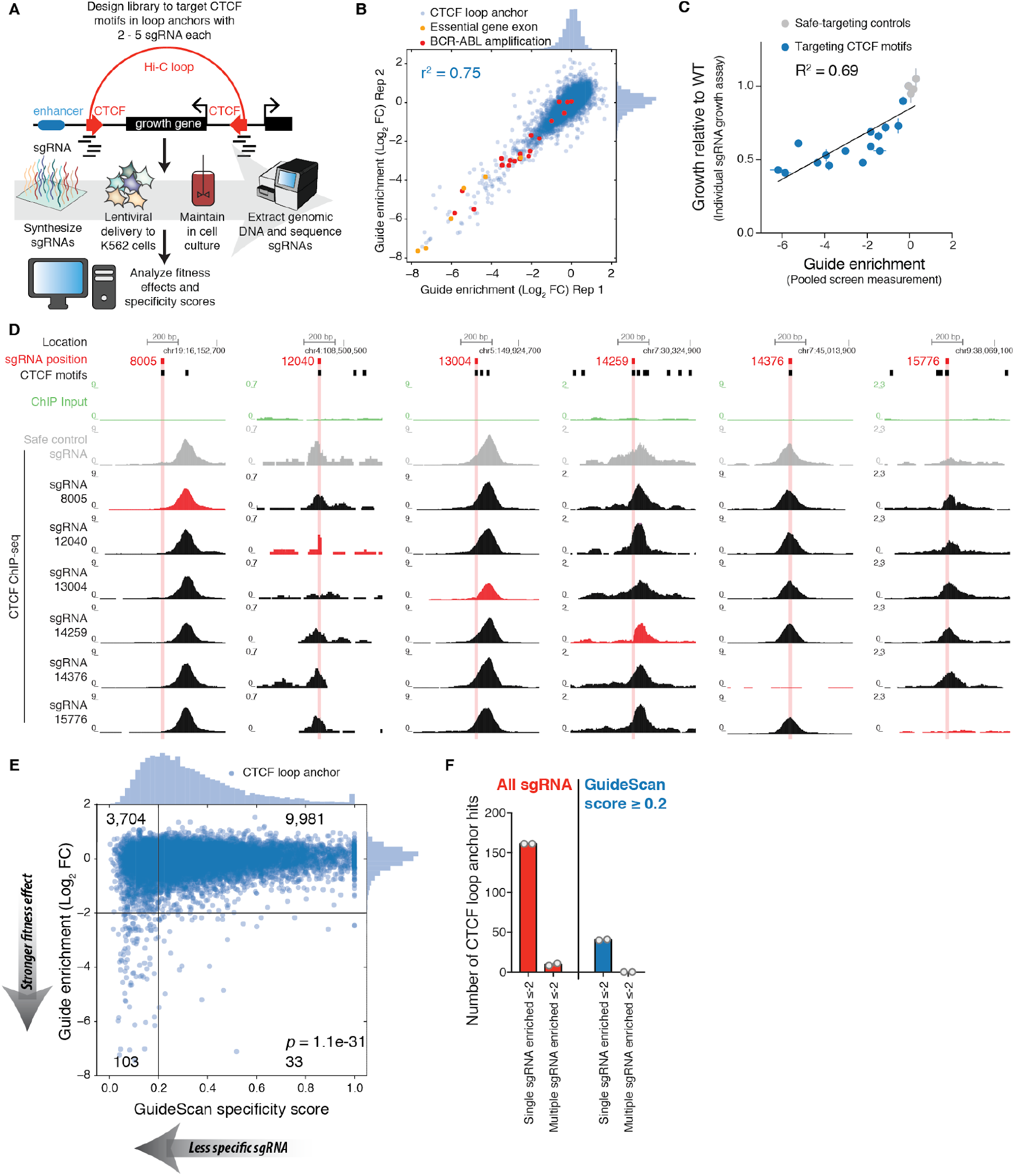
A genome-scale CRISPR-Cas9 screen finds no essential CTCF loop anchors,. A. Schematic of CTCF loop anchor motif screen, with 2 to 5 sgRNAs targeting each CTCF motif. B. Fitness effects are reproducible between independently transduced biological replicates of the screen. sgRNAs targeting essential gene exons or the BCR-ABL amplification drop out during the growth screen, as expected. Guide enrichment values are the log_2_(fold-change) of an sgRNA’s sequencing counts from after the screen compared with the original plasmid pool, computed with the casTLE screen analysis software ^7^. C. The growth effects of CTCF motif-targeting sgRNA are validated in individual competitive growth assays after lentiviral delivery of single guides to K562-Cas9 cells. Error bars are standard deviation of three technical replicates. D. CTCF ChIP-seq was performed on the K562 cells stably expressing a CTCF-targeting sgRNA. Each column presents a particular CTCF ChIP peak and the red track highlights the sgRNA that has an on-target match in that column. While some sgRNAs completely ablate CTCF binding, others only remove part of a compound CTCF ChIP peak. sgRNA 8005 targets a motif that was not in fact underlying the nearest ChIP-seq peak, likely due to problems with motif annotation or differences between K562 cell lines, yet this guide still confers a validated growth phenotype. E. Low-specificity guides are significantly enriched among CTCF motif-targeting guides with fitness effects. The Fisher’s exact test provided the p-value for the association between fitness effect and specificity using the 2 × 2 contingency table of the numbers of guides in each quadrant based on the thresholds drawn in black lines. Numbers in corners correspond to the number of CTCF site-targeting guides (blue circles) in the quadrant. The off-target search was done with GuideScan, which retrieves all off-target locations with 2 or 3 mismatches to the sgRNA spacer. sgRNAs with > 1 perfect matches to the genome or > 0 off-target locations with only 1 mismatch are not searchable within the GuideScan trie data structure and were excluded from this analysis. F. There were no CTCF motifs with concordant evidence of fitness effects from multiple high-specificity sgRNAs. Grey circles are screen biological replicates.

To better understand the mechanistic basis for these fitness effects, we characterized the transcriptional and chromatin landscape of K562 cell lines carrying mutations induced by individual sgRNAs with validated growth effects. First, we sought to confirm that sgRNAs targeting CTCF sites can disrupt CTCF binding by performing CTCF ChIP-seq on Cas9-expressing cells transduced with individual sgRNAs. Indeed, Cas9-induced indels entirely eliminated CTCF binding at 2 of the 6 motifs that we tested (Figure 1D), while they did not result in changes of CTCF occupancy at untargeted sites in the immediate vicinity or elsewhere in the genome (Supplementary Figure 1A,B). 3 of these 6 sgRNAs appeared to only partially ablate CTCF binding (in two cases likely due to the presence of other nearby CTCF motifs). A sixth sgRNA (sg8005) did not affect CTCF binding within the ChIP-seq peak, because the annotated motif we had targeted was not actually the motif underlying the peak, likely due to imperfect annotation. Surprisingly, we did not observe any changes in gene expression in the genomic neighborhoods of these motifs as measured by qPCR and RNA-seq (Supplementary Figure 1C-L). We also performed ATAC-seq for 2 of these sgRNAs and did not find significant changes in chromatin accessibility (Supplementary Figure 1M). Altogether, these data did not identify changes in gene expression or chromatin structure near the CTCF motifs as likely causes of the observed growth effects for any of the motifs we aimed to validate. Instead, we wondered whether off-target activity could explain these results, since off-target effects have previously been found to generate confounding signal in CRISPR-Cas9 growth screens ^30–32,34,35^.

### Computational model of specificity reveals major confounder in CTCF screens

To explore the possibility that off-target activity was responsible for the screen results, we retrieved specificity scores ^37^ for every sgRNA in the libraries. These sgRNA-level scores are determined by 1) searching reference genomes for off-target binding locations, 2) predicting the Cas9 activity across those sites given the pattern of mismatches between the sgRNA and the genomic DNA, and 3) aggregating these predicted Cas9 activities into a final score. Different implementations of this workflow have resulted in a variety of software tools providing specificity scores ^25,36,37,53–55^. We found that aggregate specificity scores from GuideScan ^37^ correlate well with existing data from Guide-seq ^27^, an unbiased off-target measurement assay for Cas9 (Spearman’s ρ = −0.84, Supplementary Figure 2A), so we used GuideScan scores for subsequent analyses. GuideScan scores are a weighted function of all off-target locations with 2 or 3 mismatches to the sgRNA spacer. Very low-specificity sgRNAs with > 1 perfect matches in the genome or > 0 off-target locations with only 1 mismatch are excluded from GuideScan’s trie data structure and were also excluded from our analysis.

Immediately, we observed a striking bias for low specificity scores among the sgRNAs that confer large fitness effects (*p* = 1.1e-31, Fisher’s exact test, Figure 1E). Indeed, the great majority (76%) of CTCF motif-targeting sgRNAs that have guide-level log_2_(fold-change) ≤ −2 also had GuideScan specificity scores ≤ 0.2 (on a scale of 0 to 1, where 0 indicates least specificity or greatest off-target activity), representing an 8.4-fold odds ratio. In the case of our CTCF screen, 4% of CTCF loop anchors had strong evidence of essentiality (Guide enrichment log_2_(fold-change) ≤ −2) with a single sgRNA, but only 0.2% had such evidence from multiple sgRNAs (Figure 1F). This disparity is unexpected given that the sgRNAs targeting the same site should have similar effects, but is consistent with the sgRNAs having different off-target effects. After filtering for high-specificity sgRNAs with the GuideScan score, the number of CTCF loop anchors with evidence of essentiality from multiple sgRNAs dropped to zero (out of 2,968 motifs targeted with multiple high-specificity sgRNAs).

### Fine-mapping CTCF loop anchors with Cas9

To further test whether off-target activity could explain the hits from the CTCF motif screen, we designed a fine-mapping sgRNA library targeting 270 CTCF sites, including full tilings of each such site (all possible sgRNAs within 1 kb), using up to 400 sgRNAs per site (Figure 2A). We chose CTCF sites from four categories: “hits” called by casTLE analysis before filtering with GuideScan scores, the Hi-C loop partners of these hits, non-hits, and the loop partners of the non-hits (Methods). We expected three possible results from densely tiling the loop anchors: 1) truly essential CTCF motifs would result in a strong peak of signal from high-specificity sgRNAs that generate indels near the motif (i.e. +/-20 bp), 2) regions that were essential for reasons distinct from the CTCF motif, such as being copy number amplified ^30,32,33^, would result in uniformly strong growth effects from both low- and high-specificity sgRNAs irrespective of the whether the sgRNAs overlap the motifs, and 3) non-functional motifs would only have strong signal from low-specificity sgRNAs, if any. This fine-mapping screen was performed at high coverage (~12,000 cells per sgRNA), yielded highly reproducible guide effect measurements (r^2^ = 0.92, Supplementary Figure 3A). As expected, positive control sgRNAs targeting ten essential genes were strongly depleted (Supplementary Figure 3B). We observed uniform depletion of high- and low-specificity sgRNAs tiling regions near the *BCR-ABL* amplification but not elsewhere (Supplementary Figure 3C,D), as expected. Both high- and low-specificity sgRNAs had strong growth effects when targeting exons of essential genes but no effect in the neighboring introns (Figure 2B), demonstrating that the fine-mapping screen can discern the short functionally relevant sequences of coding exons from background with high fidelity. Strikingly, the great majority (93%) of sgRNAs tiled within the 1 kb CTCF loop anchor regions and that had a strong fitness effect were, again, low-specificity guides with GuideScan scores ≤ 0.2 (p = 2.3e-233, Fisher’s exact test, Supplementary Figure 3E). While the previous motif-targeting library only used 2-5 sgRNAs per motif, this fine-mapping library included all possible guides overlapping a window of +/-20 bp of the “hit” CTCF motif centers. Despite this increase in sgRNA density, after filtering with GuideScan scores, we still found zero CTCF motifs with evidence of essentiality from multiple high-specificity sgRNAs (Figure 2C and Supplementary Figure 3F,G). We therefore concluded that the observed hits in the CTCF screens were consistent with off-target activity.

**Figure 2.**
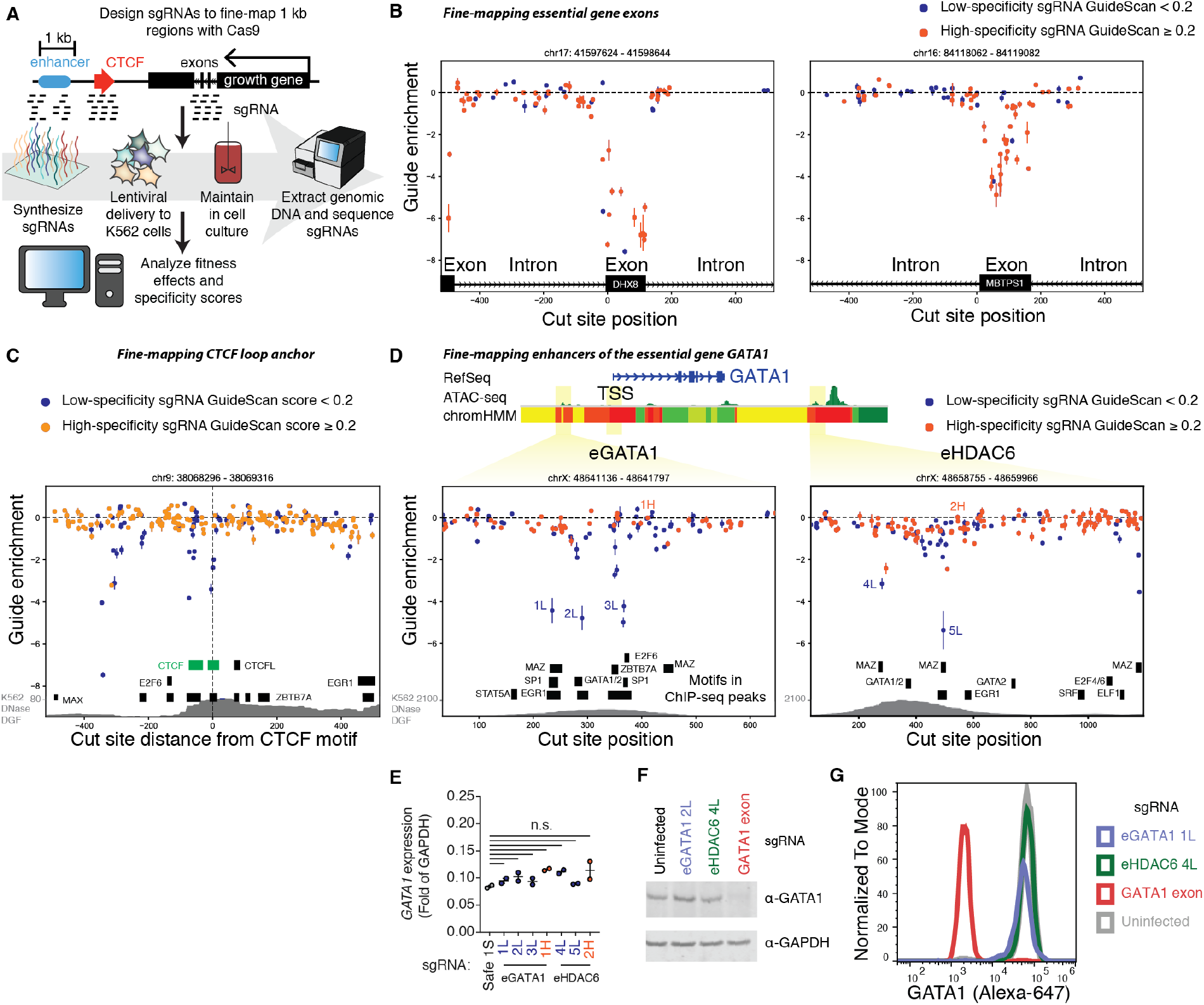
Low-specificity sgRNAs confound identification of essential motifs in fine-mapping screen of loop anchors and enhancers of essential genes. A. A fine-mapping Cas9 growth screen was performed with sgRNAs densely tiling two types of regions: 1) 1 kb windows around select hit and non-hit CTCF loop anchors from the CTCF motif screen and 2) two enhancers of *GATA1*, previously called eGATA1 and eHDAC6. B. As a positive control, we verified that the fine-mapping screen correctly maps the boundaries of exons of essential genes with high-specificity sgRNAs. Each point is the average enrichment of two biological replicates and the error bar is the standard error. C. Fine-mapping screen results from a 1 kb region centered on a motif that was a false positive hit in the original motif-targeting screen (targeted with sgRNAs 15776 and 15777 and also shown in **Figure 1** and **Supplementary Figure 1**). All evidence for the essentiality of a CTCF motif comes from low-specificity sgRNAs. Motifs in ChIP-seq peaks are shown as black boxes and CTCF motifs as green boxes. D. Fine-mapping screen results from two regions containing enhancers of the essential gene *GATA1.* sgRNAs selected for validation studies are labeled (e.g. “1L” represents the first sgRNA with a low specificity score). ChromHMM is colored according to the 15-state scheme ^56^ (briefly, reds are predicted promoter states, yellows are enhancer states, and greens are other transcriptionally active states). E. The enhancer motif-targeting sgRNAs identified in (**D**) do not significantly decrease GATA1 expression according to qPCR (p > 0.05, ANOVA). F. The sgRNAs identified in (**D**) do not significantly decrease GATA1 protein expression according to Western blot. G. The sgRNAs identified in (**D**) do not significantly decrease GATA1 protein expression according to flow cytometry for GATA1 protein level. Additional validation data are shown in **Supplementary Figure 4**.

### Off-target activity confounds identification of motifs within enhancers

To test our ability to dissect the essentiality of non-coding elements beyond chromatin loop anchors, we also fine-mapped two enhancers which regulate expression of the essential gene *GATA1* in K562 cells, tiling them with 110 and 174 sgRNAs to span the entire 611 bp and 1.1 kb regions, respectively. These enhancers, named eGATA1 and eHDAC6, were previously identified in a CRISPRi tiling growth screen in K562 ^9^, but their constituent functional motifs remain uncharacterized, a gap we sought to fill with higher resolution dissection by Cas9 fine-mapping. These screens revealed narrow peaks defined by 1-2 sgRNAs that overlapped known TF ChIP-Seq motifs within the DNase hypersensitive sites in the enhancers ^51^ (Figure 2D). However, these sgRNAs were again of low specificity, raising doubts that their targets were in fact essential motifs and motivating a careful validation of the sgRNAs and their effects on *GATA1* expression. We installed the sgRNAs individually into K562, and found that this resulted in indel mutations (37-98%) in the genomic DNA at the corresponding target motifs (Supplementary Figure 4A). These sgRNAs also caused significant growth phenotypes (Supplementary Figure 4B) which correlated with the growth effects measured in the pooled screen (r^2^ = 0.76, Supplementary Figure 4C). Strikingly, there were no concordant changes in *GATA1* expression as measured by qPCR, Western blot, or flow cytometry (Figure 2E-G and Supplementary Figure 4D). These experiments demonstrate that even sgRNAs targeting TF motifs in bona fide enhancers can have reproducible growth screen effects that are unrelated to the expression of their nearby essential gene, and that the GuideScan specificity score is useful to help identify such confounded sgRNAs.

### CRISPRi and CRISPRa off-target activity also causes confounding growth effects

CRISPRi and CRISPRa have also been used to screen for functional non-coding elements, but the potentially confounding effect of off-target activity with these platforms in the context of non-coding essential regulatory elements has not been studied. To systematically compare these technologies, we performed a tiling screen around three essential genes in K562 cells (*GATA1, MYB*, and *ZMYND8*); the library consisted of a total of 32,791 sgRNAs targeting a total of 794 kb including candidate regulatory elements, annotated exons and intervening genomic space. We screened this library with four different CRISPR-Cas9 platforms: active Cas9, nuclease-dead dCas9, CRISPRi (dCas9-KRAB ^23^), and CRISPRa (dCas9-SunTag-VP64 ^57^) (Figure 3A). As expected, in the active Cas9 screen we observed strong negative fitness effects for sgRNAs targeting exons, and in the CRISPRi screen we observe strong signals for sgRNAs targeting known essential enhancers and promoters ^9,52^ (Figure 3B and Supplementary Figure 5A-D). We also found that for CRISPRa and dCas9 screens, sgRNAs that targeted transcriptional start sites (TSS) of essential genes exhibit negative fitness effects (Figure 3B and Supplementary Figure 5D); for dCas9, this observation may be due to the binding of dCas9 interfering with the transcriptional initiation machinery ^23,58^.

**Figure 3.**
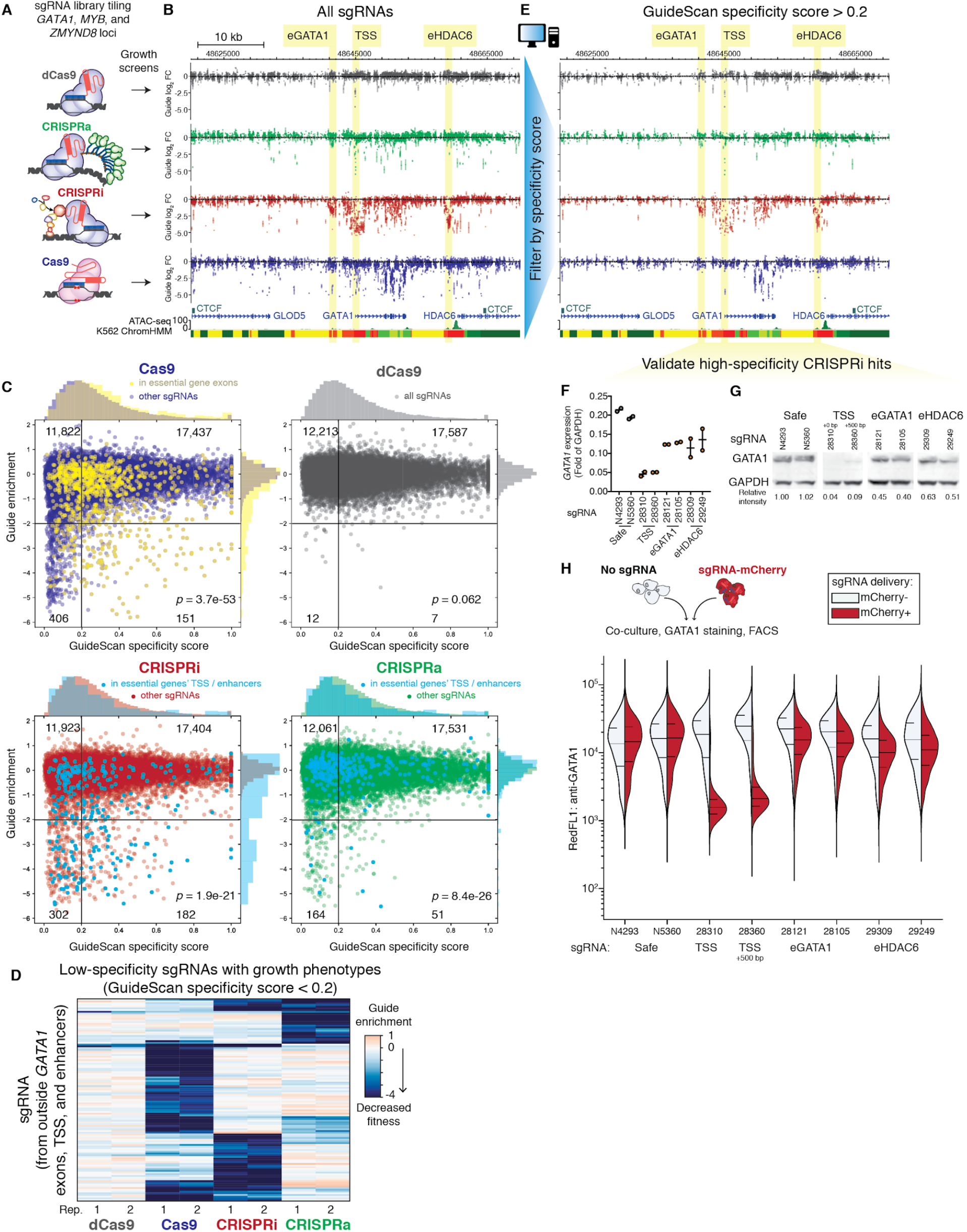
Filtering CRISPRi library for high specificity greatly reduces false positives and enables accurate detection of moderate-strength enhancers. A. Four parallel screens were conducted tiling the loci of essential growth genes *GATA1, MYB*, and *ZMYND8* using the four platforms Cas9, CRISPRa, CRISPRi and dCas9. B. Zoomed-in view of screen data around essential gene *GATA1.* Highlighted are regulatory elements with known effects on cell growth: enhancers eGATA1 and eHDAC6, and the *GATA1* transcription start site. ChromHMM is colored according to the 15-state scheme ^56^ (briefly, reds are predicted promoter states, yellows are enhancer states, and greens are other transcriptionally active states). C. Enrichment of growth effects among low-specificity sgRNAs. p-value from the Fisher’s exact test for the 2 × 2 table with quadrants as drawn and guide counts as labeled in the corners; these counts include all the sgRNAs (i.e. counts ignores the colored categories). D. Clustering of low-specificity sgRNAs reveals that each perturbation has off-target activity that reduces cell fitness with a unique subset of the low-specificity sgRNAs. Shown are the subset of sgRNAs that are upstream of eGATA1 or downstream of eHDAC6 (i.e. sgRNAs with predominantly off-target effects) and that also have a strong guide enrichment ≤ −3 in at least one replicate. E. Filtering with GuideScan specificity scores reduces noise while preserving true positive effects. F. After filtering, the CRISPRi sgRNAs in peaks have validated effects on *GATA1* expression by qPCR (p < 0.05, ANOVA). G. These CRISPRi sgRNAs also have validated effects on *GATA1* protein expression by Western blot. H. The same CRISPRi sgRNAs also have validated effects on *GATA1* protein expression by flow cytometry. Here, cells expressing an sgRNA and mCherry were co-cultured with the blank parental cell line, stained for GATA1 protein, and analyzed by flow cytometry. We then compared the distribution of GATA1 protein level between the mCherry+ and blank control cells from the same sample. Horizontal lines show the median and quartiles.

However, for each screening modality we also noticed sgRNAs with strong negative fitness effects that did not target candidate regulatory elements or annotated coding sequences and for which neighboring sgRNAs did not exhibit concordant effects (Figure 3B). Again, we suspected that the growth effects of these guides might be due to off-target activity and retrieved GuideScan specificity scores in order to investigate this possibility. Indeed, we observed a striking enrichment for low-specificity sgRNAs among the set of sgRNAs with strong negative fitness effects in the Cas9, CRISPRi, and CRISPRa screens (*p* < 1.9e-21 for all, Fisher’s exact test, Figure 3C). We questioned whether the sets of sgRNAs with putative off-target activity were highly overlapping between each CRISPR-Cas9 platform. Strikingly, this was not what we observed. In fact, sets of low-specificity sgRNAs that show significant fitness effects with Cas9, CRISPRi or CRISPRa are largely non-overlapping (Figure 3D), suggesting the off-target effects are specific to each CRISPR-Cas9 platform. Thus, off-target growth effects appear to be a function of both the sites targeted by an sgRNA and the mode of perturbation.

We questioned whether these off-target growth effects were purely a function of the absolute number of off-target sites or specific to a subset of off-target sites. We and others have shown that, in the context of coding gene screens, the number of perfect matches or 1-mismatch off-targets correlates with growth phenotypes ^34,35^. However, the analyses presented here do not include any sgRNAs with perfect genomic matches at any other place in the genome, nor sgRNAs with 1-mismatch off-targets. Across all four CRISPR-Cas9 platforms used in the tiling screens, the GuideScan score was predictive of off-target effects on cell fitness (Figure 3C and Supplementary Figure 6A), yet there was very weak correlation between growth effects and the absolute number of off-target sites (with 2 or 3 mismatches each), especially for CRISPRi/a (Supplementary Figure 6B,C). Indeed some outlier sgRNAs with thousands of off-target sites had no effects on growth. Thus, when designing and interpreting screens, the propensity to bind or cut as captured by the specificity score should be considered, rather than simply the number of off-target binding locations.

### CRISPRi screens filtered for high-specificity sgRNAs specifically detect essential regulatory elements

While the appearance of confounding off-target activity in CRISPRi screens was unexpected, GuideScan scores proved useful to identify confounded sgRNAs. We next asked if the removal of low-specificity sgRNAs would improve the reliable identification of expected regulatory elements (e.g. the TSS and the two enhancers of *GATA1*). We thus filtered out guides with GuideScan scores ≤ 0.2, which did indeed remove confounded sgRNAs while preserving strong CRISPRi signal at these enhancers and promoters (highlighted regions in Figure 3E).

To confirm that these high-specificity sgRNAs in peaks had bona fide effects on the expression of *GATA1*, we delivered single guides by lentivirus and measured *GATA1* expression by qPCR and Western blot (Figure 3F,G). Whereas targeting the *GATA1* TSS or a CRISPRi peak 500 bp downstream of the TSS both resulted in near-complete knockdown (to 4-9% of protein levels in the control cells), the enhancer-targeting sgRNAs provided partial knockdown (to 40-63% of control protein levels), and expression levels were highly correlated between RNA-level qPCR and protein-level Western blot (R^2^ = 0.92, Supplementary Figure 7A). Flow cytometry for GATA1 protein levels confirmed that CRISPRi enhancer repression resulted in partial knockdown across the population of cells, as opposed to complete silencing observed when targeting the TSS (Figure 3H). Together, these experiments validated that the high-specificity sgRNAs from the tiling CRISPRi screen resulted in on-target repression of the expected essential gene.

### CRISPRi/a off-target activity is a confounder in other non-coding growth screens

We next wondered if off-target activity might confound other CRISPRi/a non-coding growth screens for other types of elements. To directly compare the different CRISPR-Cas9 platforms with a shared library of sgRNAs, we performed parallel screens with our CTCF motif-targeting sgRNA library in K562 using CRISPRi, CRISPRa, dCas9, and Cas9 (Supplementary Figure 8A-C). When we analyzed the specificity scores of this library, we found that these CRISPRi and CRISPRa screens again showed a significant bias towards low-specificity sgRNAs having strong growth effects (Supplementary Figure 8D). The Cas9 screen in this experiment was maintained with lower coverage (cells per sgRNA) and was thus noisier than the Cas9 screen in Figure 1; interestingly, we found that this enrichment for low-specificity sgRNAs was less pronounced but remained highly significant (p = 1.1e-9, Fisher’s exact test), showing that the signature of off-target effects can be disguised in noisy screens. As with our tiling library, we found that the sets of low-specificity sgRNAs that show significant fitness effects with Cas9, CRISPRi or CRISPRa are largely non-overlapping, reproducing the previous observation that off-target effects are specific to each CRISPR-Cas9 perturbation (Supplementary Figure 8E). Again, the CRISPRi/a growth phenotypes were not reproduced when employing dCas9 with the same sgRNAs, demonstrating these off-target effects are not due to dCas9 binding alone.

To investigate the generality of these CRISPRi off-target growth effects across cell types, we retrieved GuideScan specificity scores for guide libraries from published screens targeting the promoters of genes with dCas9-KRAB-MeCP2 in SH-SY5Y and HAP1 cells ^59^. These screens found reproducible, validated hits, but also found that some sgRNAs targeting known non-essential genes had unexpected growth effects. Here, we found that these sgRNAs also had lower specificity scores (Supplementary Figure 9C). These results suggest that using CRISPRi with low-specificity sgRNAs can be associated with strong fitness effects in other cell types.

### Impact of low-specificity sgRNAs on non-coding screen designs

Finally, we investigated the extent to which non-coding elements can be targeted with high-specificity sgRNA libraries. To address this question, we characterized the distribution of GuideScan specificity scores for a number of possible screen designs. We observed that our tiling screen and CTCF site screen libraries contained significantly more low-specificity sgRNAs than Brunello ^36^, a genome-wide coding gene-targeting library (p < 0.0001, Mann-Whitney test, Figure 4A), reflecting the inherently poorer specificity of sgRNA libraries that densely tile regions or target relatively small motifs. We then designed libraries targeting all candidate cis-regulatory elements (or ccREs) which were identified in the ENCODE SCREEN databases ^60,61^. At the time of our analysis, the SCREEN databases contained 1.31 million individual ccREs, with a median length over 200 bp (Supplementary Figure 10A). We specifically focused on CRISPRi/a epigenetic perturbation designs and imposed a minimum requirement of including at least 5 sgRNAs of sufficiently high specificity for each element (to enable robust statistical analyses of functional effects at the element level). We find that 89% of SCREEN ccREs can be targeted with ≥ 5 sgRNAs at a GuideScan cutoff of 0.2 (Supplementary Figure 10B) although this varies by type of target element. For example, we find that 62% of human lncRNA TSS elements can be targeted with ≥ 5 CRISPRi sgRNAs with a specificity score > 0.2, even when selecting sgRNAs from a conservative window of only +/- 100 bp from the TSS (Figure 4B). Overall, most ccREs can be targeted with epigenome editing tools even after filtering the sgRNAs that are most likely to be confounded by off-target effects.

**Figure 4.**
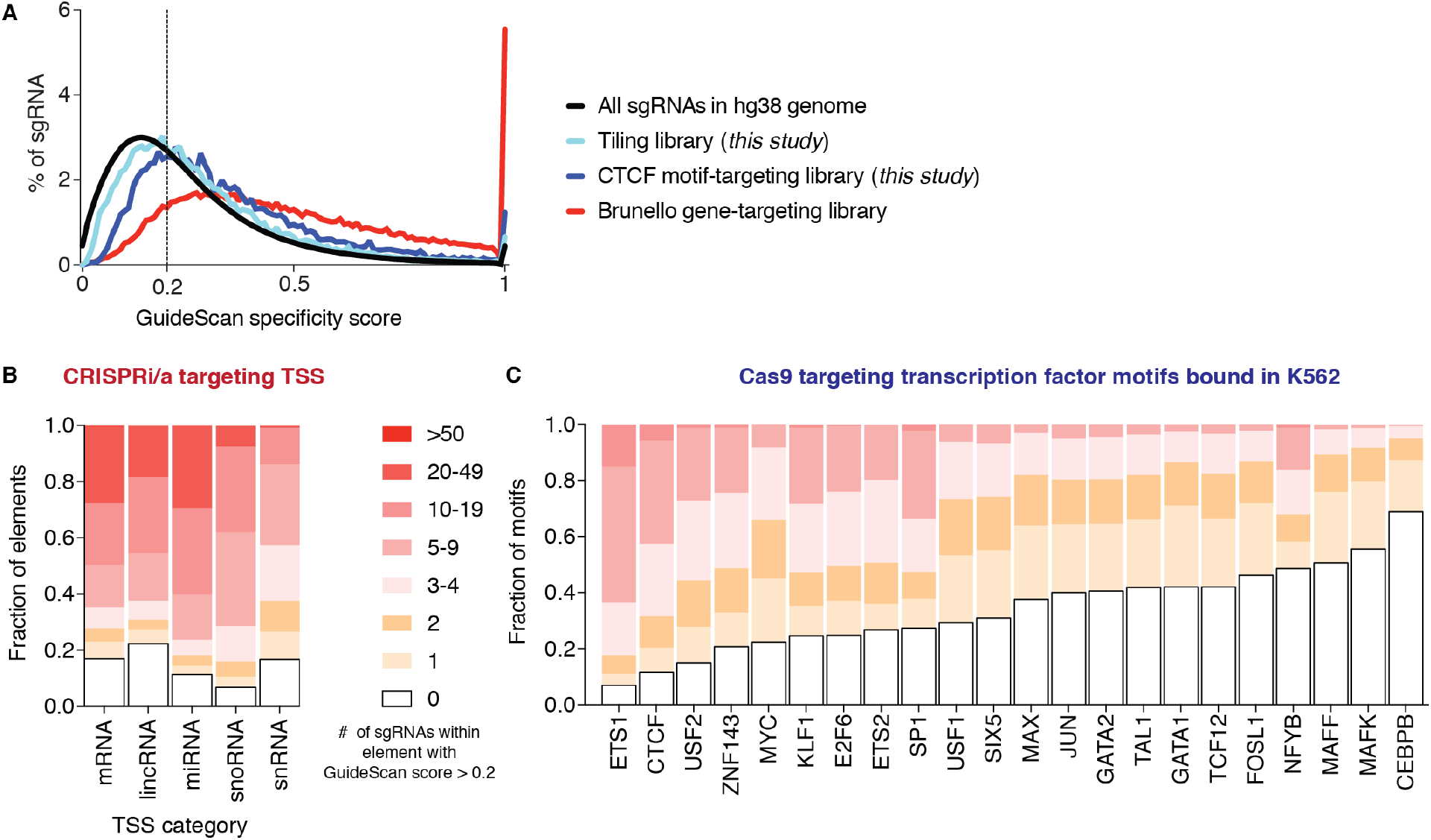
High-specificity CRISPR-Cas9 screen designs for non-coding elements. A. Distribution of GuideScan specificity scores for two non-coding libraries from this study and a gene-targeting library, in comparison to all possible sgRNA. B. Most TSSs can be targeted with multiple high-specificity sgRNA. Fraction of TSS in the ENCODE SCREEN database of ccREs that can be targeted with dCas9-based epigenome editors within a window of +/- 100bp, after filtering for GuideScan scores > 0.2. C. Fraction of motifs in TFBS motifs that can be targeted with sgRNAs with a cut site in the motif, after filtering out low-specificity sgRNAs.

However, most ccREs are composed of multiple regulatory units, such as transcription factor binding sites (TFBSs), and achieving proper mechanistic understanding of ccRE function will require perturbing these regulatory units, individually or in combination. To assess the ability of Cas9 to enable more fine-grained regulatory element mapping, we designed motif-level screens for 27 different human TFs targeting all of their annotated and occupied motifs in K562 cells and summarized the specificity score distributions for each. We find that guide specificity filtering restricts the ability to target TF motifs to a varying extent for different TFs: for example, only 31% of CEBPB motifs can be targeted with even a single overlapping sgRNA at a GuideScan cutoff of 0.2 (Figure 4C), whereas for TFs such as ETS1, 64% motifs can be targeted with 5 or more such guides. Taken as a whole, Cas9 TF motif screens, as well as splice site screens (Supplementary Figure 10C), are subject to more limiting design restrictions than screens targeting ccREs with CRISPRi/a, because the sgRNAs for these Cas9 non-coding screens must overlap the narrow target element directly while sgRNAs for CRISPRi/a ccRE screens can be selected from a larger targeting window. These designs provide a guideline for focusing future screens for essential regulatory elements on the motifs and ccREs that can be targeted with high-specificity guides.

## Discussion

Here, we found that pervasive off-target activity confounds Cas9, CRISPRi, and CRISPRa screens for essential regulatory elements by conducting several screens using sgRNA libraries designed to edit motifs and tile regions of interest in an unbiased fashion.

We and others have previously shown that off-target DNA damage from Cas9 nuclease activity affects growth screen measurements ^30–35^; this work extends these observations to non-coding growth screens. Indeed, we find that low-specificity sgRNAs are the dominant confounding factor complicating the analysis and interpretation of screens for essential regulatory elements and that, somewhat surprisingly, this conclusion holds not only for active Cas9 screens but also for dCas9-mediated perturbations such as CRISPRi and CRISPRa. Cas9 generates double-strand breaks (DSB), so a large number of off-targets for a given sgRNA could result in a major fitness effect due to cellular toxicity as a result of activation of the DNA damage response and apoptosis ^30,32–34,52^, regardless of the location of off-target sites. In contrast, dCas9-recruited epigenetic perturbations do not generate DSBs, and their off-target effects are expected to be location-dependent. Interestingly, these off-target effects cannot be fully accounted for by dCas9 binding itself, as we tested the same sgRNAs with all four CRISPR-Cas9 platforms, and nearly all sgRNAs showed reduced or unmeasurable growth effects with dCas9 alone.

As a prime example of the impact that off-target effects can have, growth screens targeting CTCF sites in K562 cells returned only hits that on closer examination were confounded by off-target activity. None of the CTCF sites that we characterized in more detail in cell lines expressing sgRNAs had a measurable impact on gene expression or chromatin states in the genomic neighborhood (Supplementary Figure 1), even when the Cas9 editing induced total loss of CTCF binding at the target motif (Figure 1D). A recent study reported that acute global degradation of all CTCF protein in cells ^50^ did not result in dramatic changes in gene expression. Thus, it is perhaps not surprising that the disruption of individual CTCF sites does not exhibit major phenotypic effects. It remains possible that some of the loop anchor CTCF motifs we targeted may be functional but redundant, or CTCF sites with the greatest functional relevance under standard growth conditions may not actually be at loop anchors. In terminally differentiated cells, such as K562, chromatin states may not be dramatically disrupted by the absence of an individual loop anchor CTCF site. The critical regulatory roles of CTCF may have to be studied in the context of embryonic development and cell differentiation, processes during which chromatin states are being established and CTCF loops likely serve an important role in the partitioning of the genome^62–65^.

Our findings have significant implications for the design and analysis of future screens. Given that 1) validation experiments of individual screen hits are time-intensive and low-throughput, and 2) there is a growing interest in global analyses of aggregated non-coding screen data, computational models for filtering out low-specificity sgRNAs are crucial to identify bona fide hits and to diagnose systemic problems before data aggregation. We find that off-target effects on cell fitness are not predictable solely from the absolute number of off-target sites for these sgRNAs, although that simple metric is often used when designing and ranking sgRNAs. In contrast, we find that the data-driven GuideScan specificity score, which accounts for the position and type of mismatches to provide a weighted assessment of Cas9’s affinity for each potential off-target site, provides a more accurate determination of off-target potential. The striking correlation of this score with fitness effects in non-coding screens, and also with direct measurements of off-target cutting using Guide-Seq, has not been described in the literature. Surprisingly, even though this score was not trained on CRISPRi/a screens, and CRISPRi/a off-targets are distinct from those of Cas9 nuclease (Figure 3D), the score was effective in identifying CRISPRi/a off-target effects.

We find that targeting a substantial fraction of individual TFBSs with high-specificity sgRNAs when using Cas9 is often impossible, although this fraction varies widely between different TFs. This constraint imposes a significant limitation on Cas9 growth screens directed at elements as small as TFBSs (< 30 bp). On the other hand, at the level of an individual ccRE (> 150 bp), sufficiently many high-specificity sgRNAs can generally be found for CRISPRi and CRISPRa screens. Notably, coding gene screens also benefit from larger available sequence from which to choose sgRNAs.

However, GuideScan models only the potential extent of off-target cleavage activity and very frequently gives low specificity scores for sgRNAs that have no effect on the phenotypic outcome of cell growth. One exciting future direction suggested by our study is the development of models to predict the phenotypic consequence of off-target activity, which can now be enabled by high-throughput datasets such as these. By integrating features including the chromatin state of off-target binding locations and the essentiality of genes near those off-target locations, it may be possible to tailor models to predict which particular sgRNAs would be confounded if used with each CRISPR-Cas9 platform.

We expect that the impact of low-specificity guides is dependent on the phenotype being screened. Low-specificity sgRNAs have a greater potential to confound growth screens, likely because proliferation is affected by many factors in the cell, while screens employing different selection strategies may be less sensitive to these effects. Studies of ccRE effects that involve measuring the RNA or protein products of cognate genes, separating cell populations according to expression levels, and then identifying the particular sgRNAs associated with each expression level may also be less affected by off-target effects. Similarly, experiments that couple CRISPR-Cas9 screens to single-cell readouts of gene expression ^66–70^ or chromatin accessibility ^71^ may likewise overcome limitations associated with growth as a readout.

Regardless, limitations remain that will be best addressed by the development of perturbation systems that either expand the targetable sequence space or minimize off-targets. Efforts in both of these directions are ongoing, e.g. devising guide design strategies that reduce off-target effects such as truncated guides ^34,72^, engineering high-specificity variants of Cas9 ^73–76^, and exploring the possibilities for adapting other CRISPR enzymes without strict PAM requirements ^16,77–79^. We expect that the combination of technological improvements, judicious screen design, and careful data analysis that explicitly considers guide specificity will enable the comprehensive functional characterization of the essential regulatory elements in the human genome.

## Materials and Methods

### Cell lines and cell culture

All experiments presented here were carried out in K562 cells (ATCC CCL-243) grown as previously described ^7^. Cells were cultured in a controlled humidified incubator at 37°C and 5% CO_2_, in RPMI 1640 (Gibco) media supplemented with 10% FBS (Hyclone), penicillin (10,000 I. U./mL), streptomycin (10,000 ug/mL), and L-glutamine (2 mM). Experiments were performed in four modified K562 cell lines: K562 stably expressing SFFV-Cas9-BFP, K562 expressing SFFV-dCas9-BFP, K562 expressing dCas9-SunTag-VP64 ^3^ (CRISPRa), and K562 expressing SFFV-dCas9-KRAB-BFP (CRISPRi). The CRISPRa cell line expressing the SunTag system was a gift from the lab of Jonathan Weissman.

### CTCF motif-targeting sgRNA library design

We selected CTCF motifs in loop anchors to target as follows. We started with 6,057 loops present in K562 cells and focused on the 4,892 loop anchors that had previously annotated motifs overlapping ChIP-seq peaks ^40^ for CTCF (using STORM ^83^), such that the CTCF motifs were convergently oriented into the loop, which is suggested to be the correct orientation for loop formation. We further restricted to 4,172 loop anchor CTCF motifs that could be targeted with with at least two sgRNAs per site, as defined by our guide filtering criteria below. Some of these targets were in exons of genes or near the BCR-ABL amplification, so they were treated separately during analysis, resulting in a final count of 4,022 “Type 0” CTCF loop anchor motifs. Finally, a set of control sgRNAs targeting safe regions was added. Briefly, safe-targeting negative control sgRNAs are highly filtered to target a non-functional genomic site and avoid having severe growth effects while controlling for the effect of inducing a double strand break (Morgens et al., 2017). An additional 310 CTCF and Rad21 sites (“Types 1 - 5”) were selected with alternative methods (Supplementary Materials & Methods) and also targeted with sgRNAs in the library, but these were filtered out during analysis and not included in Figure 1 for the sake of clarity and because this small alternative set was similarly confounded by off-target activity and lacking hits. For sites that passed our filtering criteria, we selected a maximum of 5 sgRNAs per site.

To minimize off-target effects, we filtered out sgRNAs that had exact or 1-mismatch off-target instances within a CTCF site or inside exons of GENCODEv19 ^84^ genes. We also filtered out guides with > 2 0-mismatch, > 10 1-mismatch, > 50 2-mismatch or > 200 3-mismatch genome-wide off-targets. We defined off-target matches by aligning the guides to the hg19 version of the human genome using BWA ‘aln’ with the flags -N -n 4 -o 0 -k 0 -l 7 ^85^. We also filtered out guides with too low (< 20%) or too high (> 80%) GC content and guides containing so-called “confounding oligonucleotides” that might affect the expression of the guide or PCR steps, where “confounding oligonucleotides” are defined as those that either end in “GGGGG,” contain “TTTT,” or contain restriction cut sites (“CTGCAG,” “GAAGAC,” “GTCTTC,” “CCANNNNNNTGG,” “GCTNAGC”).

### CTCF sgRNA screen execution

Oligonucleotide libraries (Supplementary Table 1) were synthesized by Agilent and then cloned into an sgRNA expression vector pMCB320 (Supplementary Table 2) that had been cut with BstXI and BlpI restriction enzymes, by ligation using T4 ligase, as previously described ^34^. Large scale lentivirus production and infection of K562-Cas9 cells were performed as previously described ^86,87^. Selection with puromycin was started three days after infection and continued for 3-4 days until the mCherry-positive percentage of cells was greater than 80%, as observed by flow cytometry on a BD Accuri. Cells were then maintained at 3,000x coverage (cells per sgRNA). Cells were maintained in log growth conditions each day by diluting cell concentrations back to a 0.5 * 10^6^ cells/mL. These conditions were also used for the dCas9, CRISPRi, and CRISPRa screens performed with this library.

Genomic DNA was extracted following Qiagen’s Blood Maxi Kit, and the guide composition was sequenced and compared to the plasmid library using casTLE ^7^ version 1.0 available at https://bitbucket.org/dmorgens/castle.

The screen was repeated in K562-Cas9 cells at 11,000x maintenance coverage for 23 days, starting from a frozen aliquot of cells after library transfection and puromycin selection (frozen at day 6). After the screen, genomic DNA was harvested and sgRNAs were amplified and sequenced as previously described ^7,88^. The high-coverage screen showed better reproducibility between biological replicates (Supplementary Figure 8C) and was used for all analyses shown in the main text (Figure 1).

### Fine-mapping screen library design

The fine-mapping screen employed densely tiled sgRNAs in short 1 kb windows around CTCF motifs, enhancers, and exons of essential genes. First, we densely tiled the regions around the CTCF motif screen hits as identified by casTLE (see below), a GC-matched set of regions around non-hit CTCFs, and the “loop partner” CTCFs that looped to any of these positive or negative CTCFs in a K562 Hi-C dataset ^40^. Non-hit CTCFs were selected from the set of CTCF sites with enrichment magnitudes less than 0.5 for all guides in all motif-targeting Cas9, CRISPRi/a, and dCas9 screens. We selected all sgRNAs provided by the GuideScan design tool within the CTCF motif and up to 500 bp on each side, for a total of 1020 bp. For each CTCF hit, we selected a 1020-bp region around a ‘GC-matched’ non-hit CTCF with a GC content within 5% of the GC content of the 1020-bp region around the CTCF hit. Additionally, we densely tiled the essential enhancers eGATA1 and eHDAC6 as positive controls and added 1000 safe-targeting guides as negative controls. As an additional positive control, we included all guides from a 10-guide gene-targeting library ^34^ for the essential genes *CTCF, RAD21, SMC1A, SMC3, MYC, GATA1, MYB, RPS28, RPS29*, and *RPS3A.*

### Fine-mapping screen execution

The screen was executed with the same protocol as the others at a maintenance coverage of approximately 12,000 K562 cells per sgRNA. After 20 days, genomic DNA was harvested and sgRNAs were amplified and sequenced with an Illumina NextSeq to a depth of 2,333 - 3,153 reads per sgRNA using a previously described protocol ^88^.

### Tiling screen library design and execution

We designed an sgRNA library (referred to from now on as the “tiling screen” library) that would allow us to compare different CRISPR-Cas9 platforms in an unbiased fashion. To this end, we decided to focus on a limited set of genes with an already known strong growth effect, specifically *GATA1* [guides covering the genomic region chrX:48544984-48752721 (in hg19 coordinates), covering a total region of 207.737 kb, with tiling density 9308/207.737kb = ~44 guides per kilobase], *MYB* (guides covering the genomic region chr6:135402680-135640267, covering a total region of 237.587 kb, with tiling density 9200/237.587kb = ~38 guides per kilobase), and *ZMYND8* (guides covering the genomic region chr20:45737857-46085556, covering a total region of 347.699 kb, with tiling density of 14282/347.699kb = ~41 guides per kilobase). These regions were determined by tiling the full annotated gene sequence and then extending the tiling for an additional 100 kb in either direction.

We filtered guides as follows. We discarded guides that had any exact or one-mismatch targets in DNase-hypersensitive sites ^60^ or exons. We also filtered out sgRNAs that had any perfect matches in the genome, or > 10 1-mismatch, > 50 2-mismatch or > 200 3-mismatch genome-wide off-targets. Matches were defined by aligning the guides to the genome using BWA ‘aln’ with the flags -N -n 4 -o 0 -k 0 -l 7 ^85^.

To allow direct comparison of effect sizes of regulatory elements in the screen with those of genes, we also included guides targeting the coding regions of the 3 genes of interest (10 guides per gene). Finally, we added a set of 1000 control guides targeting “safe” regions as defined previously ^34^.

The screen was executed with the same protocol as the others. After 14 days, genomic DNA was harvested and sgRNAs were amplified and sequenced as previously described ^88^.

### Screen data analysis

The casTLE v1.0 framework ^7^ was used to process screen data, including alignment of reads to an index of guide oligos, subsequent guide filtering, and estimation of effects on cell growth. For growth screens, enrichment scores were calculated by comparing samples from the final day (day 14, 21, or 23, depending on the screen) with the plasmid library.

For the CTCF motif screen, we ran makeIndices.py with parameters ‘-s 31 -e 37’ and makeCounts.py with parameters ‘-l 20’; we also grouped sgRNAs that target the same motif to measure motif-level effects and called hits using combined biological replicates with a 10% false discovery rate, using the script analyzeCombo.py. For the fine mapping screen, we ran makeIndices.py with parameters ‘-s −34 -e 17’ and makeCounts.py with parameters ‘-l 17 -m 0 -s -’. For the tiling screen, we ran makeIndices.py with parameters ‘-s 11 -e 17’ and makeCounts.py with parameters ‘-l 19’.

### GuideScan specificity scores

We retrieved GuideScan v1.0 ^37^ specificity scores from the webtool. GuideScan forgoes short string alignment (e.g. BWA) to find off-target locations and instead recovers locations from a pre-computed trie data structure; it then computes Cutting Frequency Determination (CFD) scores ^36^ for all off-target locations with 2 to 3 mismatches, and then aggregates them with the summation formula from the CRISPR MIT tool ^25^ (dividing 1 by the sum of 1 plus all the CFDs), such that sgRNAs with more off-target activity approach GuideScan scores of 0. GuideScan does not provide scores for sgRNAs with multiple perfect genomic matches or off-targets that only differ by 1 mismatch, which are assumed to be too poor specificity for use in experiments, so we also excluded such sgRNAs from the analyses using GuideScan.

### Competitive growth assays

Competitive growth assays were performed, similarly to a previous description ^88^}, with stable K562 lines expressing Cas9, CRISPRi, or CRISPRa that were lentivirally transduced with a vector (pMCB320) expressing the sgRNA and mCherry and then, after 2 to 3 days, selected with puromycin for 3 to 4 days, until the mCherry+ fraction of cells was > 90%. Then 40,000 of these mCherry+ cells were mixed 1:1 with blank cells from the parental line (Day 0) in 1 mL of fresh RPMI media and grown in triplicate or quadruplicate in 24-well plates. The cells were maintained at a confluence less than 1e6 cells per mL. The changes in the mCherry+ proportion of cells were measured on an Accuri BD C6 flow cytometer on Day 0, 4, and 7 and gating on mCherry expression in channel FL3.

### Motif mapping

Transcription factor motif recognition sequences were mapped genome-wide using FIMO ^89^ (version 4.12.0 of the MEME-Suite ^90^ using the CIS-BP database ^91^ as a reference set of position weight matrices.

### External datasets

Data on the fitness effect of protein coding genes in K562 cells was obtained from previously published studies ^7,52^. Uniformly processed ChIP-seq and DNAse-seq datasets were obtained from the ENCODE portal (https://encodeproject.org). Data on dCas9-KRAB-MeCP2 screens were retrieved from the published supplementary materials ^59^.

### ChromHMM annotations

ChromHMM ^56^ tracks for K562 chromatin state ^51^ were retrieved from https://egg2.wustl.edu/roadmap/data/byFileType/chromhmmSegmentations/ChmmModels/coreMarks/jointModel/final/E123_15_coreMarks_mnemonics.bed.gz and visualized with the WashU Epigenome Browser ^92^.

### ChIP-seq experiments

ChIP-seq experiments were carried out as previously described ^93^ with some modifications. Briefly, 2e7 K562 cells were pelleted at 2000 *g* for 5 minutes at 4°C and then resuspended in 1x PBS buffer; 37% formaldehyde solution (Sigma F8775) was added at a final concentration of 1%. Crosslinking was carried out at room temperature for 15 minutes, and then the reaction was quenched by adding 2.5M Glycine solution at a final concentration of 0.25M. Crosslinked cells then were pelleted 2000 *g* for 5 minutes at 4°C, washed with cold 1x PBS buffer, and stored at - 80°C.

CTCF ChIP was performed using a polyclonal anti-CTCF antibody (Millipore, 07-729). For each reaction, 100 uL of Protein A Dynabeads (Thermo Fisher 10001D) were washed 3 times with a 5 mg/mL BSA (Sigma A9418) solution. Beads were then resuspended in 1 mL BSA solution and 4 uL of CTCF antibody were added. Coupling of antibodies to beads was carried out overnight on a rotator at 4°C. Beads were again washed 3 times with BSA solution, resuspended in 100 uL of BSA solution, mixed with 900 uL sonicated chromatin and incubated overnight on a rotator at 4°C. Chromatin was sonicated using a tip sonicator (Misonix) after cells were lysed with Farnham Lysis Buffer (5 mM HEPES pH 8.0, 85 mM KCl, 0.5% IGEPAL, Roche Protease Inhibitor Cocktail), and nuclei were resuspended in RIPA buffer (1x PBS, 1% IGEPAL, 0.5% Sodium Deoxycholate, 0.1% SDS, Roche Protease Inhibitor Cocktail). The sonicated material was centrifuged at 14,000 rpm at 4°C for 15 minutes to remove cellular debris, and a portion of the supernatant was saved as input. After incubation with chromatin, beads were washed 5 times with LiCl buffer (10 mM Tris-HCl pH 7.5, 500 mM LiCl, 1% NP-40/IGEPAL, 0.5% Sodium Deoxycholate) by incubating for 10 minutes at 4°C on a rotator and then rinsed once with 1x TE buffer. Beads were then resuspended in 200 uL IP Elution Buffer (1% SDS, 0.1 M NaHCO_3_) and incubated at 65°C in a Thermomixer (Eppendorf) with interval mixing to dissociate antibodies from chromatin. Beads were separated from chromatin by centrifugation, Proteinase K was added to the supernatant and crosslinks were reversed at 65°C for ~16 hours. Input samples (100 uL) were mixed with an equal volume of IP Elution Buffer, Proteinase K was added and cross-links were reversed together with the ChIP samples. DNA was purified by phenol-chloroform-isoamyl extraction followed by MinElute column (Qiagen) clean up. DNA concentration was measured using QuBIT, and libraries were generated using the NEBNext Ultra II DNA Library Prep Kit for Illumina (NEB, E7645S). Libraries were sequenced on a NextSeq (Illumina) in a 2 × 75 bp format.

### ChIP-seq data processing

Demultipexed fastq files were initially mapped to the hg19 assembly of the human genome (female version) as 1 × 36mers using Bowtie v1.0.1 ^94^ with the following settings: ‘-v 2 -k 2 -m 1 --best --strata’, for quality assessment purposes (see AQUAS: https://github.com/kundajelab/chipseqpipeline) (Supplementary Table 3). For subsequent analyses of CTCF occupancy, reads were mapped against the female version of the hg19 assembly of the human genome using the ‘bwa mem’ algorithm in the BWA aligner with default settings and filtering non-unique and low-quality alignments using samtools ^85^ with the ‘-F 180 -q 30’ options. A consensus set of peaks was derived from the three “safe” sgRNA CTCF ChIP-seq datasets as described in the AQUAS pipeline. FRiP values ^95^ were calculated for each dataset using this set of peak calls. Read coverage tracks were generated using custom-written Python scripts. For the purpose of comparison between datasets and normalizing for differences in ChIP strength between individual experiments, tracks were rescaled as follows:

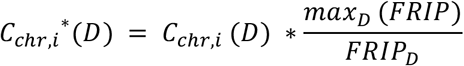

Where *C_chri,i_*(*D*) is the normalized coverage (in RPM, or Read Per Million mapped reads units) of position *i* on a given chromosome *chr* in dataset *D*, and *C_chr,i_^*^*(*D*) is the rescaled coverage.

### RNA-seq data processing and analysis

Paired-end 2 × 100 bp PolyA+ and Total RNA-seq reads were mapped using version 2.5.3a of the STAR aligner ^96^ against the hg19 version of the human genome with haplotypes removed but retaining random chromosomes, with version 19 of the GENCODE annotation ^84^ as a reference. Gene expression quantification was then carried out on the STAR alignments transformed into transcriptome space using version 1.3.0 of RSEM ^97^. Differential expression analysis was performed using DESeq2 ^98^ with the RSEM estimated read counts per gene as an input. Mapping and QC statistics are provided in Supplementary Table 4.

### ATAC-seq experiments

ATAC-seq experiments were carried out following the Omni-ATAC-seq protocol as previously described ^99^, using 50,000 K562 cells per biological replicate and two replicates per sgRNA.

### ATAC-seq analysis

Paired-end 2 × 36 bp reads were first mapped to the mitochondrial genome to assess the fraction of mitochondrial reads in each sample. All other reads were then mapped to the hg19 genome assembly using BWA as described above. Statistics are summarized in Supplementary Table 5.

### ICE analysis of indels

Cells were harvested and total genomic DNA was isolated using QuickExtract DNA Extraction Solution (VWR, Radnor, PA, cat# QE09050). PCR was prepared using 5X GoTaq Green Reaction Buffer and GoTaq DNA Polymerase (Promega, Madison, WI, cat# M3005), 10 mM dNTPs, and primers designed approximately 250-350 basepairs upstream and 450-600 basepairs downstream of the predicted cut site. PCR reactions were run on a C1000 Touch Thermo Cycler (Bio-Rad). PCR products were then purified over an Econospin DNA column (Epoch, Missouri City, TX, cat# 1910-250) using Buffers PB and PE (Qiagen, Hilden, Germany, cat# 19066 and cat# 19065). Sanger sequencing ab1 data were obtained from Quintara Biosciences and editing efficiency of knockout cell lines were analyzed using Synthego’s online ICE Analysis Tool (https://ice.synthego.com) ^81^.

### RT-qPCR experiments

RNA from 100,000 K562 cells was extracted with RNA QuickExtract (Lucigen QER090150). RNA was treated with DNaseI from the same kit, reverse transcribed with AMV RT (Sigma 10109118001), and then cDNA were quantified in multiplex TaqMan qPCR reactions using commercially available probe sets (Thermo Fisher 4453320) and TaqMan FastAdvanced Master mix (Thermo Fisher 4444556). 3 to 4 technical qPCR replicates were used for each biological replicate.

### Flow cytometry for GATA1 protein levels

We devised a flow cytometry assay wherein we co-culture cells expressing the sgRNA and mCherry from a lentivirus with non-transduced cells and stain for *GATA1* protein. Staining of GATA1 protein levels was performed as previously described ^100^. Specifically, cells were fixed with Fix Buffer I (BD Biosciences) for 15 minutes at 37°C. Cells were washed with 10% FBS in PBS once and then permeabilized on ice for 30 min using Perm Buffer III (BD Biosciences). Cells were washed twice and then stained with anti-GATA1 primary (1:1000, rabbit, Cell Signalling Technologies cat no. 3535S) for 1 hour at 4°C. After two more washes, cells were incubated with Goat anti-rabbit antibody conjugated to Alexa Fluor 647 (1:1000, ThermoFisher cat no. A-21244) for 1 hour at 4°C. After a final round of washing, flow cytometry was performed using a FACScan flow cytometer (BD Biosciences). We analyzed the data with CytoFlow by gating the cells on mCherry expression and then plot the *GATA1* protein level in mCherry+ and non-transduced cells. This approach controls for variability in staining efficiency as the two cell groups are mixed within the same sample.

### Western blot for GATA1 protein levels

Cells transduced with a lentiviral vector containing an sgRNA and puromycin-T2A-mCherry were selected with puromycin (1μg/mL) were selected until mCherry was > 85%. 1 million cells were lysed in lysis buffer (1% Triton X-100, 150mM NaCl, 50mM Tris pH 7.5, 1mM EDTA, Protease inhibitor cocktail). Protein amounts were quantified using the DC Protein Assay kit (Bio-Rad). Equal amounts were loaded onto a gel and transferred to a nitrocellulose membrane. Membrane was probed using GATA1 antibody (1:1000, rabbit, Cell Signalling Technologies cat no. 3535S) and GAPDH antibody (1:2000, mouse, ThermoFisher cat no. AM4300) as primary antibodies. Donkey anti-rabbit IRDye 680 LT and goat anti-mouse IRDye 800CW (1:20,000 dilution, LI-COR Biosciences, cat nos. 926-68023 and 926-32210, respectively) were used as secondary antibodies. Blots were imaged on a LiCor Odyssey CLx.

## Data availability

We will submit the following datasets to accessible online repositories: CRISPR-Cas9 screen data (tiling screens, fine-mapping screen, CTCF motif screens), CTCF ChIP-seq.

## Acknowledgments

We thank Evan Boyle, Maxwell Mumbach, Avanti Shrikumar, Kyuho Han, and Nasa Sinnott-Armstrong for helpful conversations and assistance. We thank Christina Leslie, Yuri Pritykin, Andrea Ventura and other members of the Leslie lab for helpful conversations about GuideScan. We thank the Stanford Functional Genomics Facility for sequencing ATAC-seq libraries. J.T. is supported by the NSF GRFP. M.C.B. is supported by a grant from Stanford ChEM-H and an NIH Director’s New Innovator Award (1DP2HD08406901). O.U. is supported by a Howard Hughes Medical Institute International Student Research Fellowship and a Gabilan Stanford Graduate Fellowship. D.H.P. was supported by NIGMS and NHGRI of the NIH under award numbers R35GM128645 and R00HG008662 respectively. This work was supported by a grant from NIH/ENCODE 5UM1HG009436-02 to W.J.G., A.K., and M.C.B.

## Author contributions

M.W. and O.U. designed sgRNA libraries with assistance from J.T., D.M., I.M.K., P.G.G., D.H.P., and M.C.B. J.T., G.K.M, G.T.H., B.K.E., A.T., A. and A.E.T. performed experiments. J.T., M.W., G.K.M., O.U. and G.T.H. analyzed data with assistance from D.M., I.M.K., L.B., W.J.G., A.K., and M.C.B. G.K.M. analyzed scores for guides targeting motifs and ENCODE SCREEN elements. D.Y., K.S., A.L., and A.T. generated sgRNA libraries. J.T., M.W., and G.K.M. wrote the manuscript with contributions from all authors. M.P.S., L.B., W.J.G., A.K., and M.C.B. supervised the project.

## Supplementary Figures

**Supplementary Figure 1.**
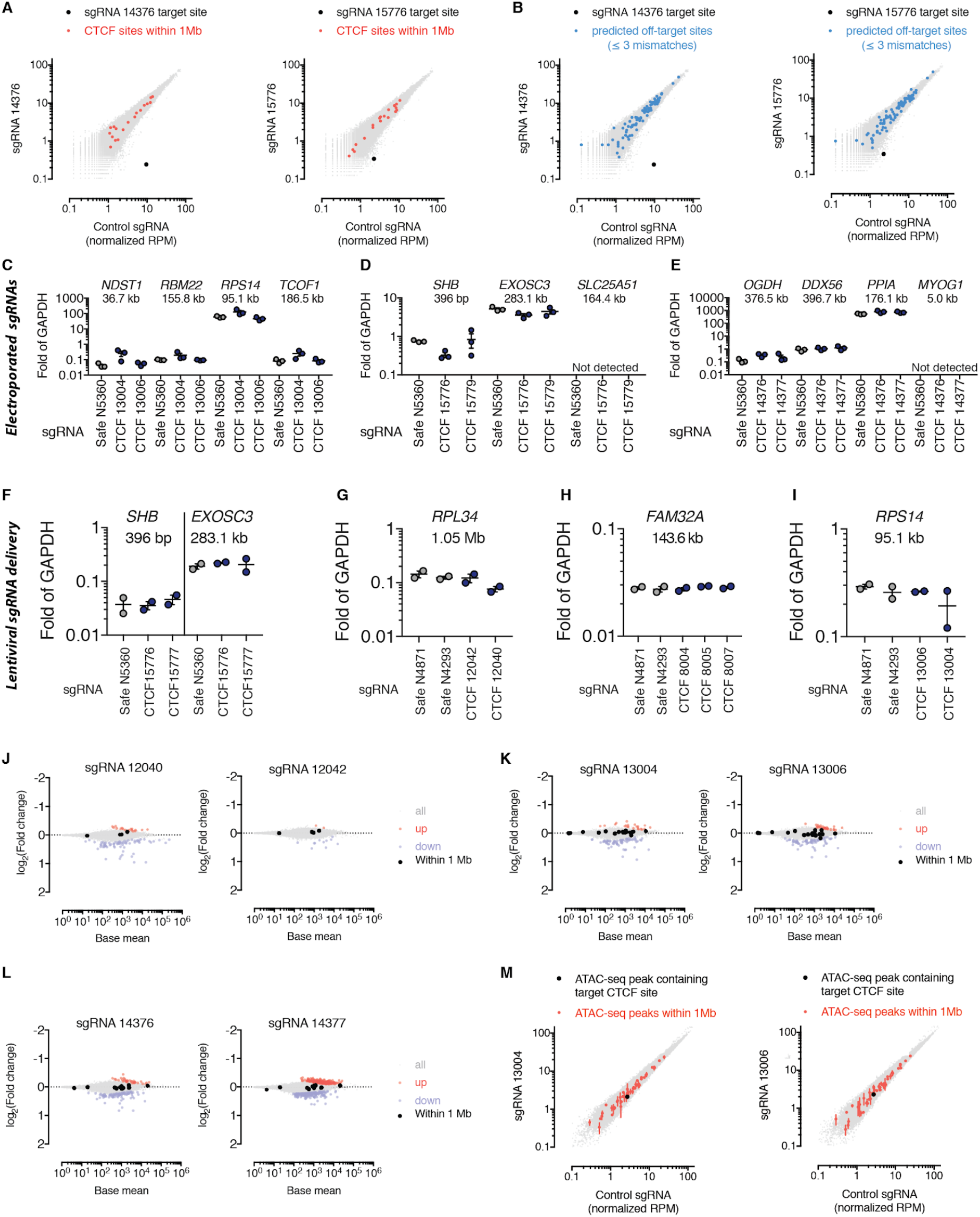
Follow-up studies of individual sgRNAs targeting CTCF motifs. A. sgRNA targeting CTCF sites were delivered via lentivirus to a K562-Cas9 cell line and then CTCF ChIP-seq was performed. No other CTCF peaks within 1 Mb of the on-target location were significantly affected. B. No other CTCF peaks that overlap a predicted off-target site with ≤ 3 mismatches were affected. List of off-target sites was provided by the Cas OFFinder webtool ^80^. C. No significant changes in the expression of nearby essential genes were detected for any of the CTCF-targeting sgRNA that were individually tested. sgRNA-mCherry plasmids were delivered by electroporation, 36 hours later the cells were confirmed to be > 70% mCherry+ by flow cytometry and RNA was extracted for qPCR. *NDST1* is a non-essential gene and the CTCF motif falls within one of its introns. *RBM22, RPS14*, and *TCOF1* are the nearest essential genes. The distances shown below the gene names are between the CTCF motif and the TSS of the gene. D. *SHB* is a non-essential gene and the CTCF motif falls within its 5’ UTR; *EXOSC3* and *SLC25A51* are the nearest essential genes. E. *MYOG1* is a non-essential gene and the CTCF motif falls within its intron. *OGDH, DDX56, and PPIA* are the nearest essential genes. Genes are determined to be essential if they were called as hits with a 10% FDR in previous Cas9 ^34^, or CRISPRi/a gene screens ^52^. F. Individual sgRNAs were delivered by lentivirus, 2 days later cells were selected for sgRNA delivery with puromycin, and 5 days after delivery RNA was extracted for qPCR. Both sgRNAs labeled “CTCF” (i.e. sgRNAs 15776 and 15777) target the same CTCF motif. Same target motif as in **D**. G. *RPL34* is the nearest essential gene. H. *FAM32A* is the nearest essential gene. I. *RPS14* is the nearest essential gene.. J. The lenti-transduced cells were subjected to RNA-seq and the mRNA expression fold-changes compared to safe-targeting sgRNAs is shown. The two sgRNAs target the same CTCF motif. None of the black dots (genes within 1 Mb of the motif) are significantly differentially expressed. K. As in **J** for another target CTCF motif. L. As in **J** for another target CTCF motif. M. No changes in ATAC-seq peaks in the cells stably expressing CTCF-targeting sgRNAs 13004 or 13006.

**Supplementary Figure 2.**
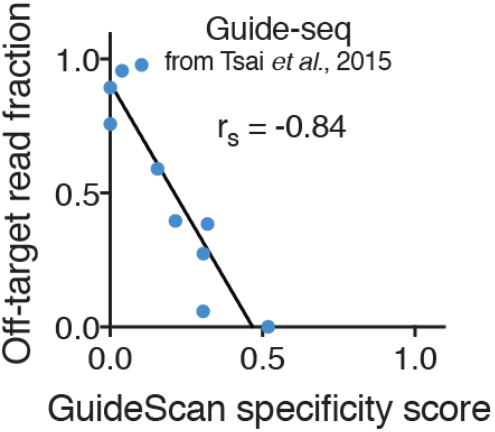
Validation of guide-level GuideScan specificity scores with an unbiased off-target assay. We retrieved GuideScan specificity scores for sgRNAs that were tested for off-target cleavage with the unbiased, genome-wide assay Guide-seq ^27^. The scores correlate with the off-target read fraction, defined as the fraction of total Guide-seq reads that align to off-target sites. Some sgRNAs did not have GuideScan scores because they had multiple perfect genomic matches or off-targets with only 1 mismatch; these sgRNAs were given a score of 0 for this analysis.

**Supplementary Figure 3.**
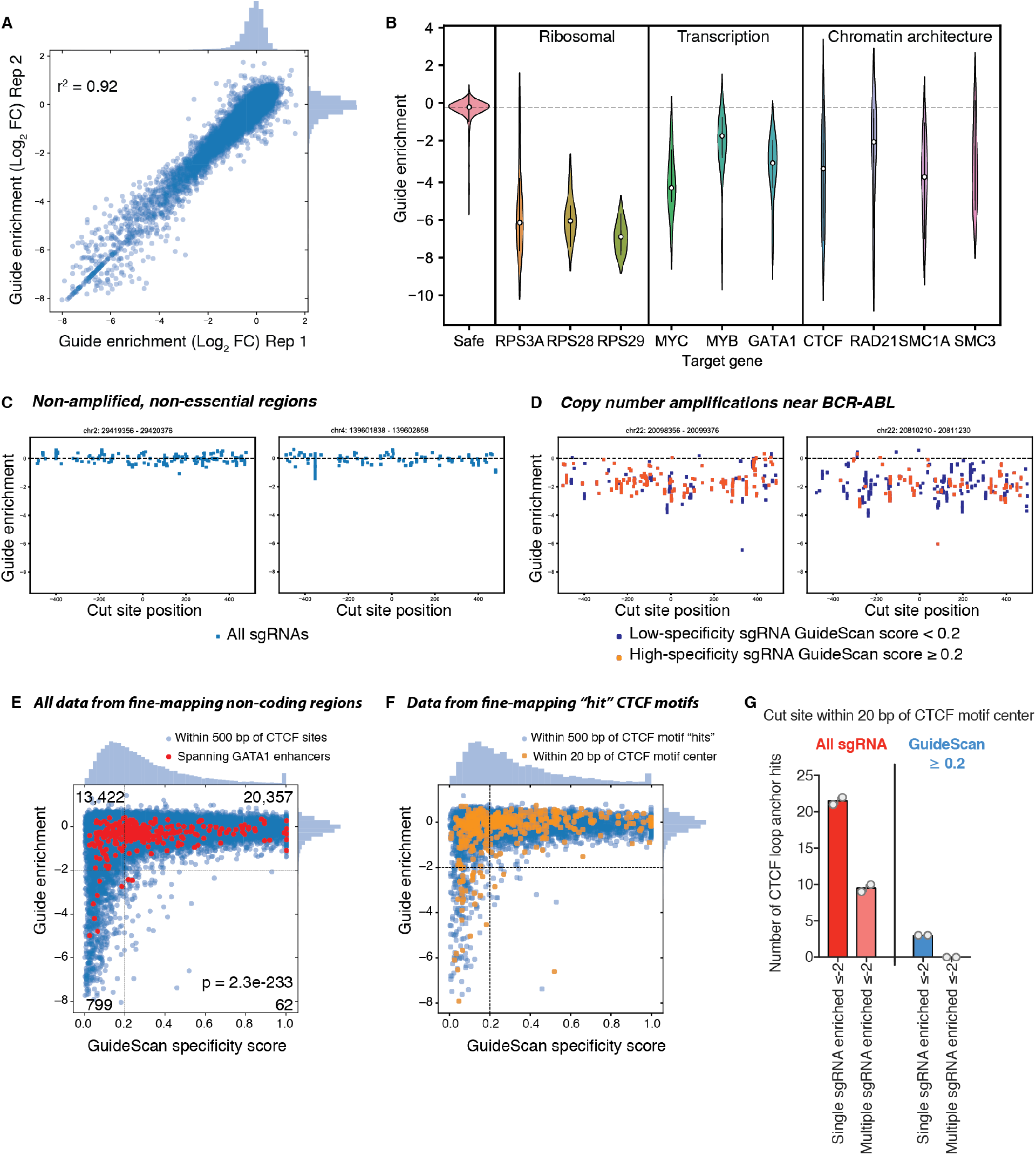
Fine-mapping screen confirms confounding effect of off-target activity. A. Reproducibility of biological replicates from a growth screen using the fine-mapping library. B. Positive controls demonstrate successful detection of essential genes. The targeted genes are essential ^7^, meaning that targeting them should decrease cell growth. Each gene was targeted with 10 sgRNAs in its coding regions; the distribution of sgRNAs is shown, and the functional annotation of each gene is labeled. “Safe” refers to safe-targeting negative control sgRNA. C. Examples of two non-amplified regions without any essential elements or any sgRNA confounded by off-target activity. D. Examples of two copy number amplified regions near *BCR-ABL* showing a distinct uniform depletion that is unrelated to the specificity of the sgRNAs. E. Low-specificity sgRNAs, in both the CTCF-anchor and GATA1-enhancer regions, are significantly enriched to have growth effects (p-value from Fisher’s exact test). F. Shown is the subset of the fine-mapping screen from 1 kb windows around motifs that previously had evidence of strong essentiality in the CTCF motif-targeting screen. G. There were no CTCF motifs with concordant evidence of fitness effects from multiple high-specificity sgRNAs, despite targeting 37 CTCF motifs with multiple high-specificity sgRNA and these CTCF sites previously being called as “hits” in the CTCF motif-targeting sgRNA screen. Grey circles are screen biological replicates and the bar marks the mean value.

**Supplementary Figure 4.**
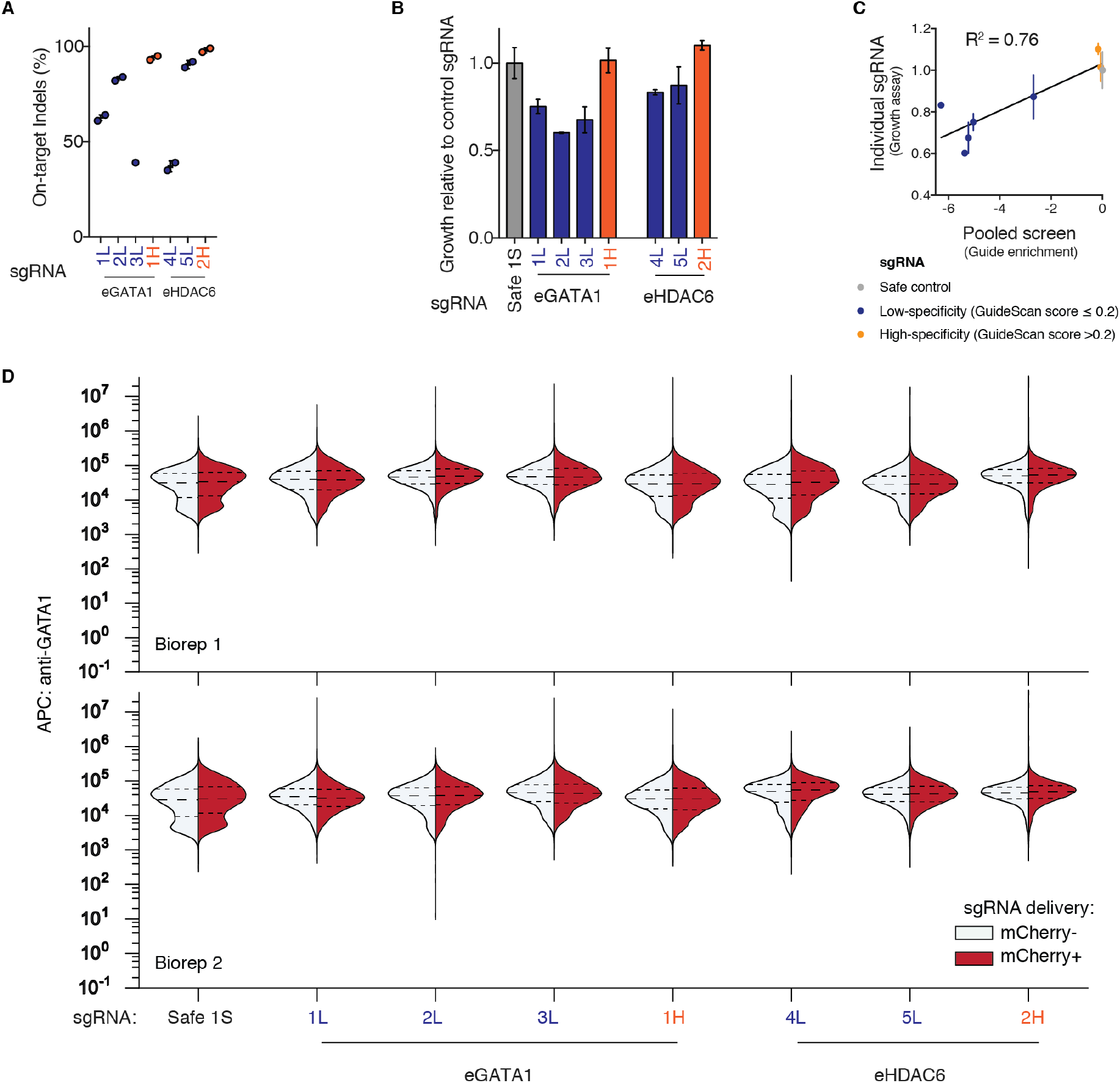
Validation experiments for fine-mapping screen of enhancers of *GATA1.* A. Individual sgRNAs generated on-target indels in K562 after lentiviral delivery and puromycin selection, as quantified by ICE analysis ^81,82^. B. Competitive growth assay validated expected growth effects in these individual cell lines. C. Individually measured growth effects correlate with the pooled screen measurements. D. Additional flow cytometry for GATA1 protein levels confirmed there was no change in expression of GATA1 in these cell lines. Cells transduced with the sgRNA-mCherry lentiviral vector were co-cultured with non-transduced parental cells and then stained and analyzed by FACS together in order to control for variation in staining efficiency between samples. In all samples, the distribution of GATA1 levels is not significantly different between the mCherry+ and blank cells. Dashed lines within the histograms mark the quartiles. sgRNA labeled as in Figure 2.

**Supplementary Figure 5.**
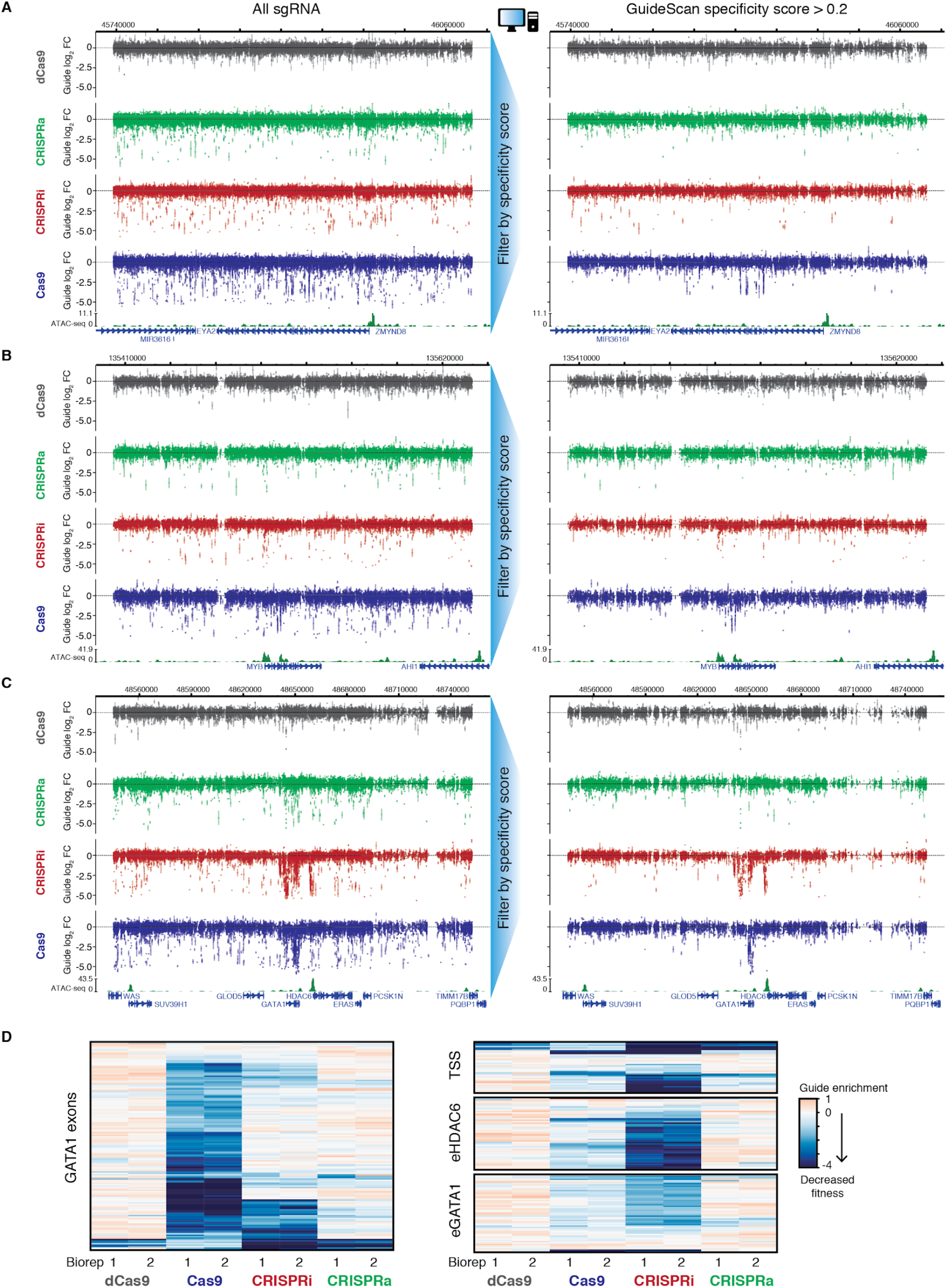
Tiling screens of three regions around essential genes with four CRISPR-Cas9 perturbations. A. Four parallel screens were conducted tiling the loci of essential growth genes *GATA1, MYB*, and *ZMYND8* using the four platforms Cas9, CRISPRa, CRISPRi and dCas9. Shown is the full tiled region around *ZMYND8* with and without filtering for high-specificity sgRNAs with the GuideScan score. B. Full tiled region around *MYB.* C. Full tiled region around *GATA1.* D. Clustering of sgRNAs from the *GATA1* tiling screen that target regions with expected on-target effects (exons, TSS, and enhancers).

**Supplementary Figure 6.**
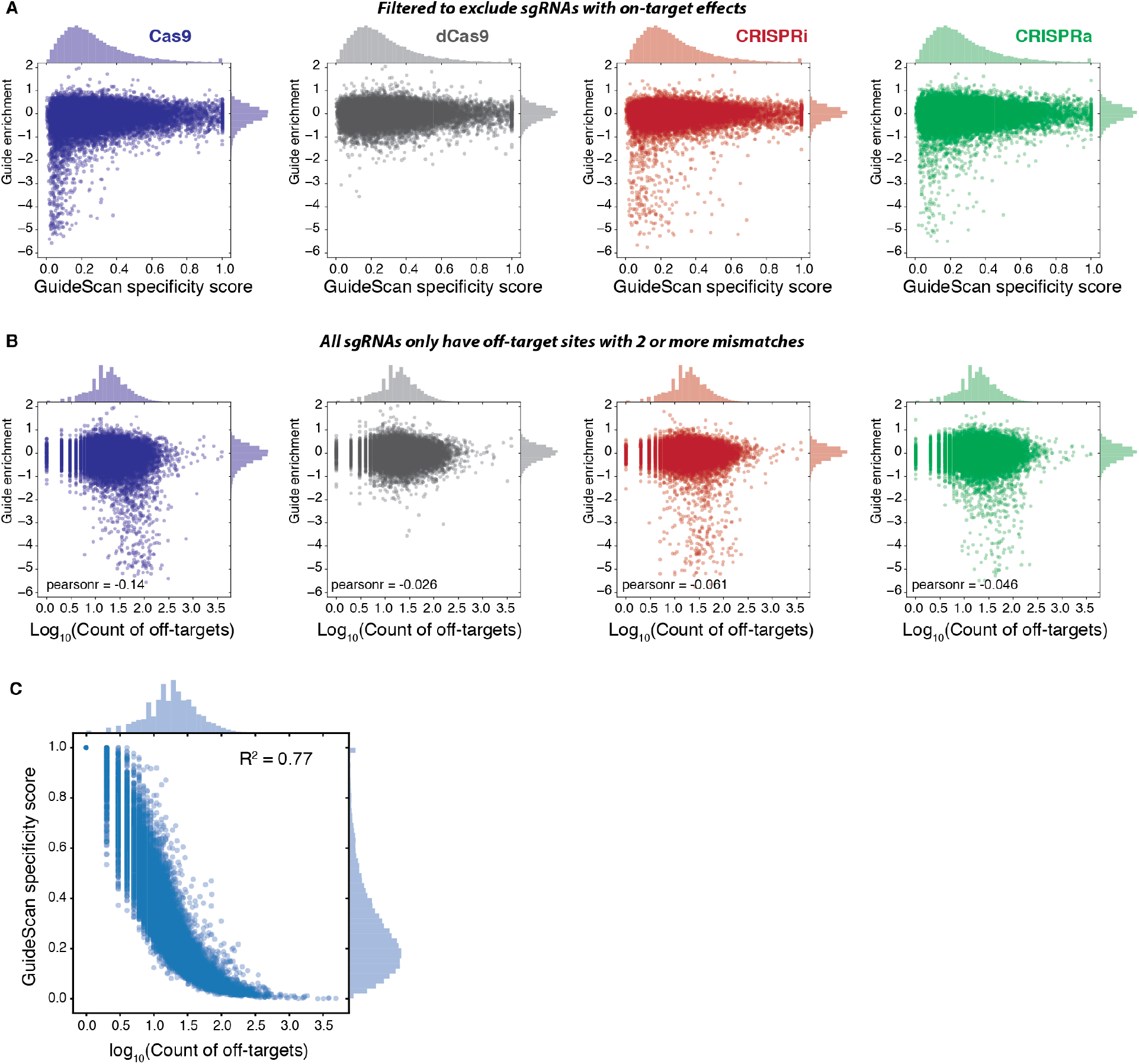
Comparison of fitness effects and specificity scores with the number of off-target binding locations. A. Comparison of GuideScan scores with fitness effects in the tiling screen, filtered to exclude sgRNAs that are likely to have on-target growth effects by removing sgRNAs 1000 bp upstream to 1000 bp downstream of *ZMYND8* or *MYB* coding sequences, and 1000 bp upstream of eGATA1 to 1000 bp downstream of eHDAC6. For the similar plot that includes those sgRNAs, see Figure 3C. sgRNAs with multiple perfect matches to the genome or off-target locations with only 1 mismatch are not searchable within the GuideScan trie data structure and were excluded from this library. B. For the same set of sgRNAs in **A**, we compared the guide enrichment from the tiling screen with the number of off-target binding locations that have 2-3 mismatches. The off-target search was done with GuideScan. C. For comparison, the relationship between the GuideScan specificity score and the number of off-target locations for the same sgRNAs in the tiling screen library.

**Supplementary Figure 7.**
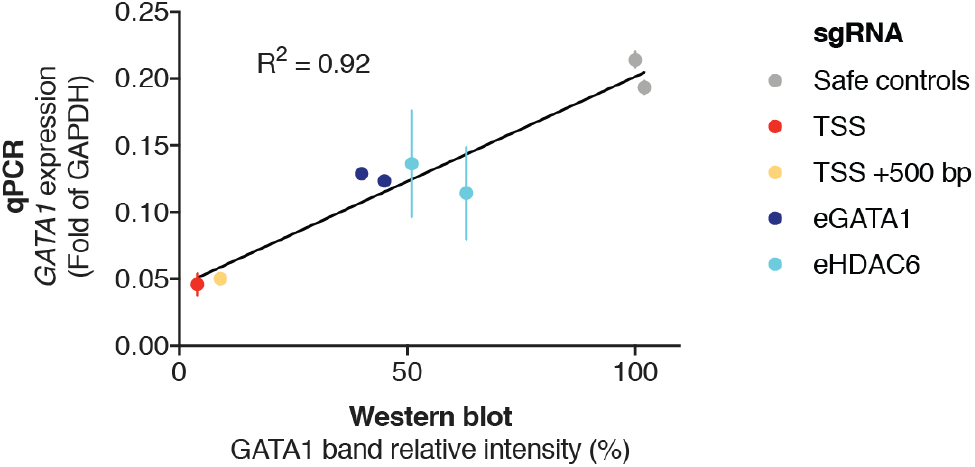
Validation of CRISPRi repression of essential enhancers with high-specificity sgRNAs. After delivery of individual sgRNA by lentivirus, followed by puromycin selection, we performed qPCR for GATA1 mRNA levels and a Western blot for GATA1 protein levels (shown in Figure 3). The knockdown measurements are correlated.

**Supplementary Figure 8.**
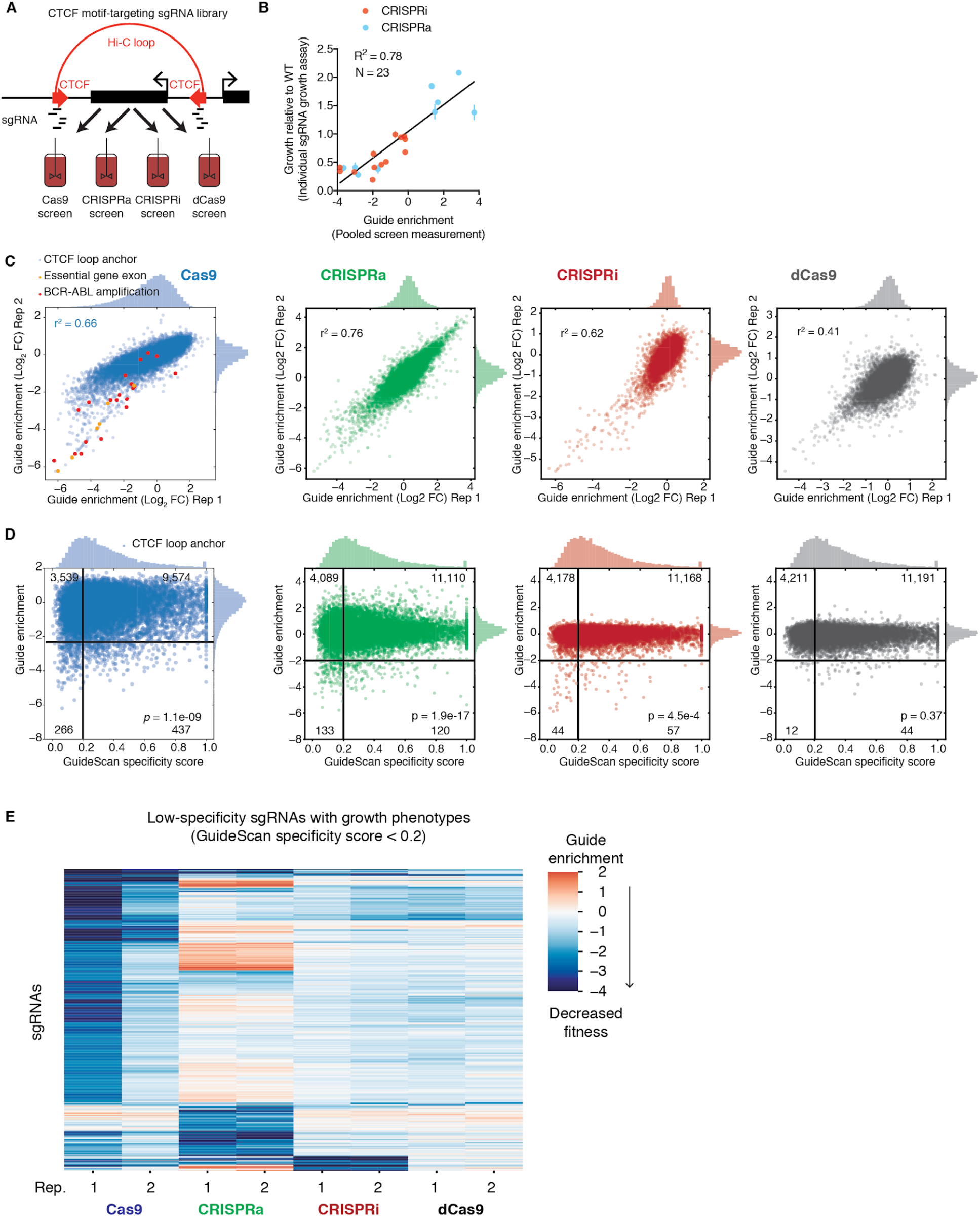
Parallel screens of CTCF loop anchors with Cas9, CRISPRi/a, and dCas9. A. The CTCF motif-targeting sgRNA library was used in parallel screens to compare the CRISPR-Cas9 platforms. All screens shown here were maintained at 3000x coverage (cells per sgRNA), whereas the Cas9 screens shown in Figure 1 were maintained at 11,000x coverage. B. Growth effects measured in this screen were validated with individual competitive growth assays. Validation of Cas9 effects shown in Figure 1. Error bars are standard deviation of three technical replicates. C. Reproducibility between biological replicates. For CRISPRi/a, sgRNAs ≤1000 bp from the TSS of an essential gene identified in a previous CRISPRi/a gene screen were excluded to avoid on-target artifacts. D. Low-specificity guides are significantly enriched among CTCF motif-targeting guides with fitness effects when using CRISPRi/a. P-value from Fisher’s exact test, using a 2 × 2 table of the numbers of guides in each quadrant based on the thresholds drawn in black lines. Numbers in corners correspond to the number of CTCF site-targeting guides in the quadrant. sgRNAs with > 1 perfect matches to the genome or > 0 off-target locations with only 1 mismatch were excluded from this analysis, as before. Notably, the Cas9 screen shown here was maintained at lower coverage and thus resulted in noisier data than the replicates shown in Figure 1. It showed a significant, but less pronounced, enrichment for low-specificity guides among the guides with fitness effects (Fisher’s exact test) than in the higher quality screen data shown in Figure 1, showing that experimental noise can disguise the confounding effect of off-target activity. E. Clustering of low-specificity sgRNAs reveals that each perturbation has off-target activity that reduces cell fitness with a unique subset of the low-specificity sgRNAs. Shown are the subset of low-specificity sgRNAs that have a guide enrichment ≤−2 in at least one replicate.

**Supplementary Figure 9.**
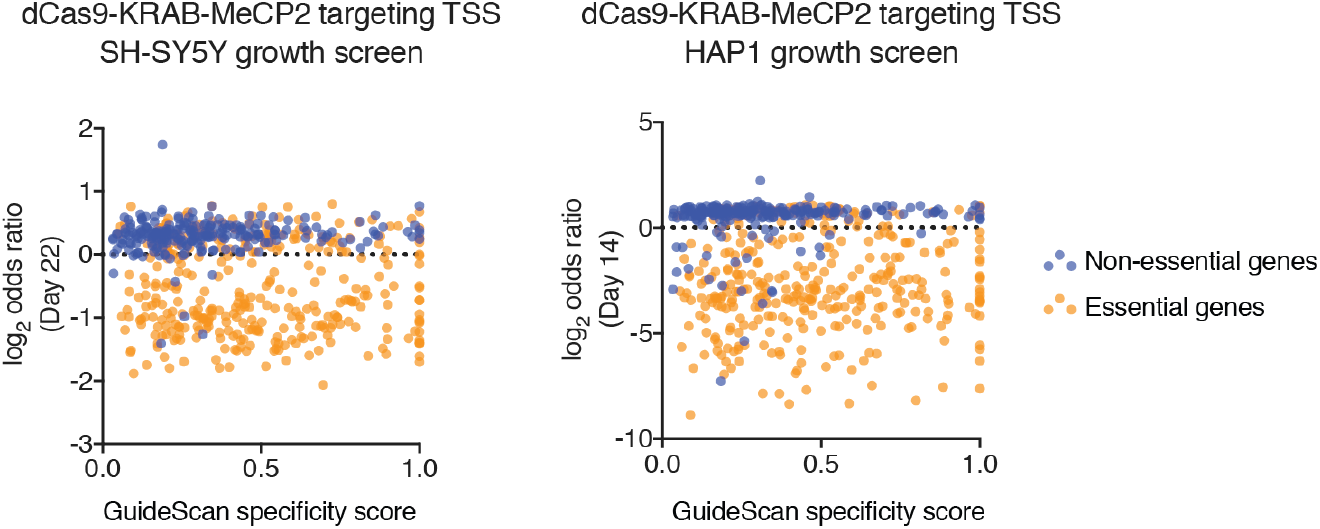
Low-specificity sgRNAs can have growth effects in other cell types with other forms of CRISPRi. We retrieved data from a published growth screen where sgRNAs were targeted to the TSS of known essential and non-essential genes ^59^, in different cell types. The marked depletion of sgRNAs targeting non-essential genes was unexpected and the authors discussed the need for further investigations to clarify the source of these effects. Here, we found that these sgRNAs have low specificity scores, implicating off-target activity. However, the enrichment was not significant, possibly due to the small number of sgRNAs in the dataset.

**Supplementary Figure 10.**
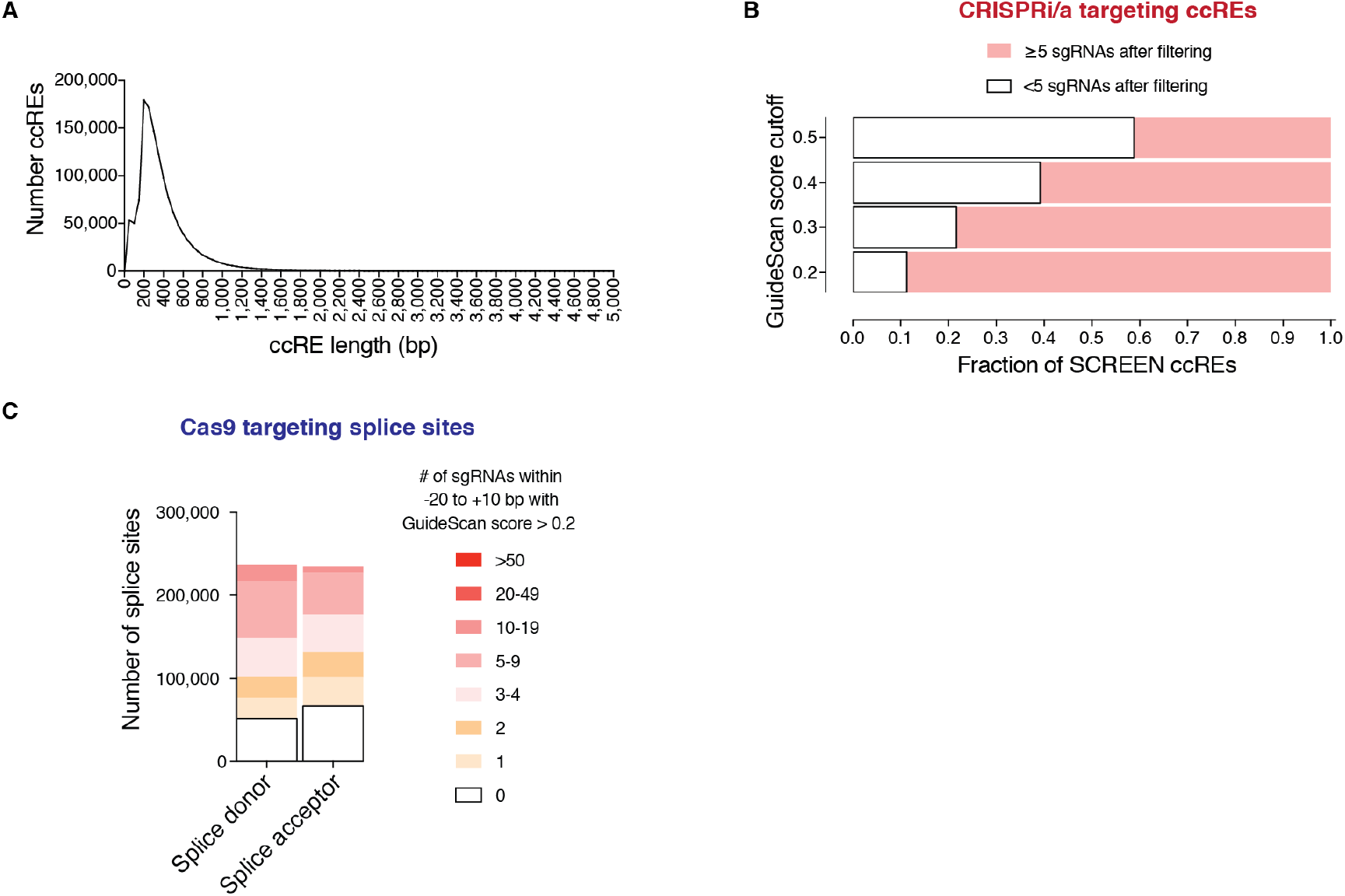
Filtered library designs for regulatory elements and splice sites. A. ccREs were retrieved from the ENCODE SCREEN database and their distribution of lengths is shown. B. Various GuideScan score filtering cutoffs were applied to the sets of sgRNAs overlapping the ccREs. 89% of ccREs can be targeted with ≥5 sgRNAs with GuideScan scores > 0.2, enabling CRISPRi/a screens of ccREs with high-specificity libraries. C. Fraction of splice sites that can be targeted with sgRNAs within a window (−20 to +10 bp), after filtering out low-specificity sgRNAs.

