## Supplementary Information for "Identification and mitigation of pervasive off-target activity in CRISPR-Cas9 screens for essential non-coding elements"

### Supplementary Tables

#### Supplementary Table 1: sgRNA libraries used in this study

The sgRNA sequences and the sgRNA scores as retrieved from GuideScan are provided in a separate Excel file.

#### Supplementary Table 2: sgRNA sequences, primers, and plasmids used in this study

| | |
|---|---|
| sgRNA N4871 (Safe - Cas9, CRISPRi, CRISPRa) | GGAAAATGATGGTCTGCAAC |
| sgRNA N5360 (Safe - Cas9, CRISPRi, CRISPRa) | GCACATTTGGATTTCATGTC |
| sgRNA N4293 (Safe - Cas9, CRISPRi, CRISPRa) | GAGGAGAGCCAATGATCTCT |
| sgRNA N5284 (Safe - Cas9, CRISPRa) | GTGTCCTTGTTTAGAAAGCA |
| sgRNA 13004 (CTCF - Cas9) | GAGAGGGGGCCTCCAGAGGG |
| sgRNA 13006 (CTCF - Cas9) | GGAGCAGAGGGGGCCTCCAG |
| sgRNA 15776 (CTCF - Cas9, CRISPRi) | GAGCTGCCGGCAGGAGGCGG |
| sgRNA 15777 (CTCF - Cas9, CRISPRi) | GGAGAGCTGCCGGCAGGAGG |
| sgRNA 15779 (CTCF - Cas9) | GGAGCCAGAGAGCTGCCGGC |
| sgRNA 14376 (CTCF - Cas9) | GTTCCCAGGGGCTCCCACCA |
| sgRNA 14377 (CTCF - Cas9) | GTCCCCCTGGTGGGAGCCCC |
| sgRNA 12040 (CTCF - Cas9) | GCCCACCAGGGAGCAGCATG |
| sgRNA 12042 (CTCF - Cas9) | GGCCCCATGCTGCTCCCTGG |
| sgRNA 8004 (CTCF - Cas9, CRISPRi) | GCTAGCCAAAGGACCAGGAG |
| sgRNA 8005 (CTCF - Cas9, CRISPRi) | GTAGCCAAAGGACCAGGAGA |
| sgRNA 8007 (CTCF - Cas9, CRISPRi) | GGCCAAAGGACCAGGAGAGG |
| sgRNA 14259 (CTCF - Cas9) | GCGTGGGAGCCGGAGGATGG |
| sgRNA 14261 (CTCF - Cas9) | GGGGAGCCGGAGGATGGCGG |
| sgRNA 15923 (CTCF - CRISPRi) | GTGGTTGAGGGACCAGGAGG |
| sgRNA 15926 (CTCF - CRISPRi) | GTTGTGGTTGAGGGACCAGG |

| | |
|---|---|
| sgRNA 15699 (CTCF - CRISPRi) | GTATTCTAGCACTTGCCCAC |
| sgRNA 15703 (CTCF - CRISPRi) | GGCTCATTGGCTCCACCCAG |
| sgRNA 5636 (CTCF - CRISPRi) | GACGTTCCCACTCTCCCTCC |
| sgRNA 5635 (CTCF - CRISPRi) | GGCACCACCTGGAGGGAGAG |
| sgRNA 15171 (CTCF - CRISPRa) | GAGTGACTGTCCTTCCACCA |
| sgRNA 15173 (CTCF - CRISPRa) | GTGACTGTCCTTCCACCAGG |
| sgRNA 13189 (CTCF - CRISPRa) | GTGTGGACGTGAGGGGGCAC |
| sgRNA 13190 (CTCF - CRISPRa) | GGCAGAATGTGGACGTGAGG |
| sgRNA 7138 (CTCF - CRISPRa) | GGTGAGCACCAGGAGGAGGG |
| sgRNA 7140 (CTCF - CRISPRa) | GGTGTGAGCACCAGGAGGAG |
| sgRNA 11698 (CTCF - CRISPRa) | GAGGACTCCAGGGCCCACAG |
| sgRNA 11699 (CTCF - CRISPRa) | GCTCCAGGGCCCACAGAGGG |
| sgRNA 16209 (CTCF - CRISPRa) | GGGCCCCTGGAGGCAGGAGT |
| sgRNA 16210 (CTCF - CRISPRa) | GGCCCAACTCCTGCCTCCAG |
| sgRNA 1S (Safe) | GCACATTTGGATTTCATGTC |
| sgRNA 1L (eGATA1) | GTTGGGGGAGACGAGGGCGG |
| sgRNA 2L (eGATA1) | GCAAGGAGGCAGCTGGGAGT |
| sgRNA 3L (eGATA1) | GACGGGGATGGGGGAGGGAA |
| sgRNA 1H (eGATA1) | GCGGGGTTTCCAGCTCTTGC |
| sgRNA 4L (eHDAC6) | GGCGGCAGGACATCTTCAAG |
| sgRNA 5L (eHDAC6) | GGGGAGTTGCGGGGGAGAGG |
| sgRNA 2H (eHDAC6) | GACACTTTCTATTACTGCTT |
| sgRNA N4293 (Safe) | GAGGAGAGCCAATGATCTCT |
| sgRNA 28310 (TSS) | GGTGATCCCAGGGGGTGTCC |
| sgRNA 28360 (TSS +500 bp) | GTAGAGCAGATAAGGGGTTT |
| sgRNA 28121 (eGATA1) | GGCGCCCTACTCCTCACCCT |
| sgRNA 28105 (eGATA1) | GGAATCAGTGGGGCCAGATC |
| sgRNA 28309 (eHDAC6) | GTGTGGCTGGGCCGAGAGCG |
| sgRNA 29249 (eHDAC6) | GAGATAGGTTCACGTAGGAG |

| | |
|---|---|
| Fwd primer ICE (eGATA1) | GGCAAACGCTTGACTCCTTA |
| Rev primer ICE (eGATA1) | CTCCTTGTGGTGCAAGTGTG |
| Fwd primer ICE (eHDAC6) | GCCTGCAAAGCCTAAGATGT |
| Rev primer ICE (eHDAC6) | AGGCATCGTTGTAAACCACA |
| sgRNA-mCherry lentiviral plasmid pMCB320 (https://www.addgene.org/89359/) | gcttaagcggtcgacggatcgggagatctcccgatcccctatggtgcactctcagtacaatctgctctgatgcc<br>gcatagttaagccagtatctgctccctgcttgtgtgttggaggtcgctgagtagtgcgcgagcaaaatttaagc<br>tacaacaaggcaaggcttgaccgacaattgcatgaagaatctgcttagggttaggcgttttgcgctgcttcgcg<br>atgtacgggccagatatacgcgttgacattgattattgactagttattaatagtaatcaattacggggtcatta<br>gttcatagcccatatatggagttccgcgttacataacttacggtaaatggcccgcctggctgaccgcccaacga<br>cccccgcccattgacgtcaataatgacgtatgttcccatagtaacgccaatagggactttccattgacgtcaat<br>gggtggagtatttacggtaaactgcccacttggcagtacatcaagtgtatcatatgccaagtacgccccctatt<br>gacgtcaatgacggtaaatggcccgcctggcattatgcccagtacatgaccttatgggactttcctacttggca<br>gtacatctacgtattagtcatcgctattaccatggtgatgcggttttggcagtacatcaatgggcgtggatagc<br>ggtttgactcacggggatttccaagtctccaccccattgacgtcaatgggagtttgttttggcaccaaaatcaa<br>cgggactttccaaaatgtcgtaacaactccgccccattgacgcaaatgggcggtaggcgtgtacggtgggaggt<br>ctatataagcagcgcgttttgcctgtactgggtctctctggttagaccagatctgagcctgggagctctctggc<br>taactagggaacccactgcttaagcctcaataaagcttgccttgagtgcttcaagtagtgtgtgcccgtctgtt<br>gtgtgactctggtaactagagatccctcagacccttttagtcagtgtggaaaatctctagcagtggcgcccgaa<br>cagggacttgaaagcgaaagggaaaccagaggagctctctcgacgcaggactcggcttgctgaagcgcgcacgg<br>caagaggcgaggggcggcgactggtgagtacgccaaaaattttgactagcggaggctagaaggagagagatggg<br>tgcgagagcgtcagtattaagcgggggagaattagatcgcgatgggaaaaaattcggttaaggccagggggaaa<br>gaaaaaatataaattaaaacatatagtatgggcaagcagggagctagaacgattcgcagttaatcctggcctgt<br>tagaaacatcagaaggctgtagacaaatactgggacagctacaaccatcccttcagacaggatcagaagaactt<br>agatcattatataatacagtagcaaccctctattgtgtgcatcaaaggatagagataaaagacaccaaggaagc<br>tttagacaagatagaggaagagcaaaacaaaagtaagaccaccgcacagcaagcggccggccgcgctgatcttc<br>agacctggaggaggagatatgagggacaattggagaagtgaattatataaatataaagtagtaaaaattgaacc<br>attaggagtagcacccaccaaggcaaagagaagagtggtgcagagagaaaaaagagcagtgggaataggagctt<br>tgttccttgggttcttgggagcagcaggaagcactatgggcgcagcgtcaatgacgctgacggtacaggccaga<br>caattattgtctggtatagtgcagcagcagaacaatttgctgagggctattgaggcgcaacagcatctgttgca<br>actcacagtctggggcatcaagcagctccaggcaagaatcctggctgtggaaagatacctaaaggatcaacagc<br>tcctggggatttggggttgctctggaaaactcatttgcaccactgctgtgccttggaatgctagttggagtaat<br>aaatctctggaacagatttggaatcacacgacctggatggagtgggacagagaaattaacaattacacaagctt<br>aatacactccttaattgaagaatcgcaaaaccagcaagaaaagaatgaacaagaattattggaattagataaat<br>gggcaagtttgtggaattggtttaacataacaaattggctgtggtatataaaattattcataatgatagtagga<br>ggcttggtaggtttaagaatagtttttgctgtactttctatagtgaatagagttaggcagggatattcaccatt<br>atcgtttcagacccacctcccaaccccgaggggacccgacaggcccgaaggaatagaagaagaaggtggagaga<br>gagacagagacagatccattcgattagtgaacggatcggcactgcgtgcgccaattctgcagacaaatggcagt<br>attcatccacaattttaaaagaaaaggggggattggggggtacagtgcaggggaaagaatagtagacataatag<br>caacagacatacaaactaaagaattacaaaaacaaattacaaaaattcaaaattttcgggtttattacagggac<br>agcagagatccagtttggttagtaccgggcccgctctagagatccgacgcgccatctctaggcccgcgccggcc<br>ccctcgcacggacttgtgggagaagctcggctactcccctgccccggttaatttgcatataatatttcctagta<br>actatagaggcttaatgtgcgataaaagacagataatctgttctttttaatactagctacattttacatgatag<br>gcttggatttctataacttcgtatagcatacattatacgaagttataaacagcacaaaaggaaactcaccctaa<br>ctgtaaagtaattgtgtgttttgagactataaGtatcccttggagaaCCAcctTGTTGGACCAGGATGGGCACC<br>ACCCGTTTAAGAGCTAAGCTGGAAACAGCATAGCAAGTTTAAATAAGGCTAGTCCGTTATCAACTTGAAAAAGT<br>GGCACCGAGTCGGTGCTTTTTTTCtcgagtactaggatccattaggcggccgcgtggataaccgtattaccgcc<br>atgcatGTGCCCGTCAGTGGGCAGAGCGCACATCGCCCACAGTCCCCGAGAAGTTGGGGGGAGGGGTCGGCAAT<br>TGAACCGGTGCCTAGAGAAGGTGGCGCGGGGTAAACTGGGAAAGTGATGTCGTGTACTGGCTCCGCCTTTTTCC<br>CGAGGGTGGGGGAGAACCGTATATAAGTGCAGTAGTCGCCGTGAACGTTCTTTTTCGCAACGGGTTTGCCGCCA<br>GAACACAGGTAAGTGCCGTGTGTGGTTCCCGCGGGCCTGGCCTCTTTACGGGTTATGGCCCTTGCGTGCCTTGA<br>ATTACTTCCACCTGGCTGCAGTACGTGATTCTTGATCCCGAGCTTCGGGTTGGAAGTGGGTGGGAGAGTTCGAG<br>GCCTTGCGCTTAAGGAGCCCCTTCGCCTCGTGCTTGAGTTGAGGCCTGGCCTGGGCGCTGGGGCCGCCGCGTGC<br>GAATCTGGTGGCACCTTCGCGCCTGTCTCGCTGCTTTCGATAAGTCTCTAGCCATTTAAAATTTTTGATGACCT<br>GCTGCGACGCTTTTTTTCTGGCAAGATAGTCTTGTAAATGCGGGCCAAGATCTGCACACTGGTATTTCGGTTTT<br>TGGGGCCGCGGGCGGCGACGGGGCCCGTGCGTCCCAGCGCACATGTTCGGCGAGGCGGGGCCTGCGAGCGCGGC<br>CACCGAGAATCGGACGGGGGTAGTCTCAAGCTGGCCGGCCTGCTCTGGTGCCTGGCCTCGCGCCGCCGTGTATC<br>GCCCCGCCCTGGGCGGCAAGGCTGGCCCGGTCGGCACCAGTTGCGTGAGCGGAAAGATGGCCGCTTCCCGGCCC<br>TGCTGCAGGGAGCTCAAAATGGAGGACGCGGCGCTCGGGAGAGCGGGCGGGTGAGTCACCCACACAAAGGAAAA<br>GGGCCTTTCCGTCCTCAGCCGTCGCTTCATGTGACTCCACGGAGTACCGGGCGCCGTCCAGGCACCTCGATTAG<br>TTCTCGcGCTTTTGGAGTACGTCGTCTTTAGGTTGGGGGGAGGGGTTTTATGCGATGGAGTTTCCCCACACTGA<br>GTGGGTGGAGACTGAAGTTAGGCCAGCTTGGCACTTGATGTAATTCTCCTTGGAATTTGCCCTTTTTGAGTTTG<br>GATCTTGGTTCATTCTCAAGCCTCAGACAGTGGTTCAAAGTTTTTTTCTTCCATTTCAGGTGTCGTGAgCTAGC<br>CTAGCCCACCATGACCGAGTACAAGCCCACGGTGCGCCTCGCCACCCGCGACGACGTCCCCCGGGCCGTACGCA<br>CCCTCGCCGCCGCGTTCGCCGACTACCCCGCCACGCGCCACACCGTCGACCCGGACCGCCACATCGAGCGGGTC<br>ACCGAGCTGCAAGAACTCTTCCTCACGCGCGTCGGGCTCGACATCGGCAAGGTGTGGGTCGCGGACGACGGCGC<br>CGCGGTGGCGGTCTGGACCACGCCGGAGAGCGTCGAAGCGGGGGCGGTGTTCGCCGAGATCGGCCCGCGCATGG<br>CCGAGTTGAGCGGTTCCCGGCTGGCCGCGCAGCAACAGATGGAAGGCCTCCTGGCGCCGCACCGGCCCAAGGAG<br>CCCGCGTGGTTCCTGGCCACCGTCGGCGTCTCGCCCGACCACCAGGGCAAGGGTCTGGGCAGCGCCGTCGTGCT<br>CCCCGGAGTGGAGGCGGCCGAGCGCGCCGGGGTGCCCGCCTTCCTGGAGACCTCCGCGCCCCGCAACCTCCCCT<br>TCTACGAGCGGCTCGGCTTCACCGTCACCGCCGACGTCGAGGTGCCCGAAGGACCGCGCACCTGGTGCATGACC<br>CGCAAGCCCGGTGCCGGATCGGGAGAGGGCAGAGGAAGTCTGCTAACATGCGGTGACGTCGAGGAGAATCCTGG<br>CCCAccggtcgccaccatggtgagcaagggcgaggaggataacatggccatcatcaaggagttcatgcgcttca<br>aggtgcacatggagggctccgtgaacggccacgagttcgagatcgagggcgagggcgagggccgcccctacgag<br>ggcacccagaccgccaagctgaaggtgaccaagggtggccccctgcccttcgcctgggacatcctgtcccctca<br>gttcatgtacggctccaaggcctacgtgaagcaccccgccgacatccccgactacttgaagctgtccttccccg<br>agggcttcaagtgggagcgcgtgatgaacttcgaggacggcggcgtggtgaccgtgacccaggactcctccctg<br>caggacggcgagttcatctacaaggtgaagctgcgcggcaccaacttcccctccgacggccccgtaatgcagaa<br>gaagaccatgggctgggaggcctcctccgagcggatgtaccccgaggacggcgccctgaagggcgagatcaagc<br>agaggctgaagctgaaggacggcggccactacgacgctgaggtcaagaccacctacaaggccaagaagcccgtg<br>cagctgcccggcgcctacaacgtcaacatcaagttggacatcacctcccacaacgaggactacaccatcgtgga<br>acagtacgaacgcgccgagggccgccactccaccggcggcatggacgagctgtacaagtaagagaattcgtcga<br>gggacctaataacttcgtatagcatacattatacgaagttatacatgtttaagggttccggttccactaggtac<br>aattcgatatcaagcttatcgataatcaacctctggattacaaaatttgtgaaagattgactggtattcttaac<br>tatgttgctccttttacgctatgtggatacgctgctttaatgcctttgtatcatgctattgcttcccgtatggc<br>tttcattttctcctccttgtataaatcctggttgctgtctctttatgaggagttgtggcccgttgtcaggcaac |

| | |
|---|---|
| | gtggcgtggtgtgcactgtgtttgctgacgcaacccccactggttggggcattgccaccacctgtcagctcctt<br>tccgggactttcgctttcccctccctattgccacggcggaactcatcgccgcctgccttgcccgctgctggac<br>aggggctcggctgttgggcactgacaattccgtggtgttgtcggggaaatcatcgtcctttccttggctgctcg<br>cctgtgttgccacctggattctgcgcgggacgtccttctgctacgtcccttcggccctcaatccagcggacctt<br>ccttcccgcggcctgctgccggctctgcggcctcttccgcgtcttcgccttcgccctcagacgagtcggatctc<br>cctttgggccgcctccccgcatcgataccgtcgacctcgatcgagacctagaaaaacatggagcaatcacaagt<br>agcaatacagcagctaccaatgctgattgtgcctggctagaagcacaagaggaggaggaggtgggttttccagt<br>cacacctcaggtacctttaagaccaatgacttacaaggcagctgtagatcttagccactttttaaaagaaaagg<br>ggggactggaagggctaattcactcccaacgaagacaagatatccttgatctgtggatctaccacacacaaggc<br>tacttccctgattggcagaactacacaccagggccagggatcagatatccactgacctttggatggtgctacaa<br>gctagtaccagttgagcaagagaaggtagaagaagccaatgaaggagagaacacccgcttgttacaccctgtga<br>gcctgcatgggatggatgacccggagagagaagtattagagtggaggtttgacagccgcctagcatttcatcac<br>atggcccgagagctgcatccggactgtactgggtctctctggttagaccagatctgagcctgggagctctctgg<br>ctaactagggaacccactgcttaagcctcaataaagcttgccttgagtgcttcaagtagtgtgtgcccgtctgt<br>tgtgtgactctggtaactagagatccctcagacccttttagtcagtgtggaaaatctctagcagcatgtgagca<br>aaaggccagcaaaaggccaggaaccgtaaaaaggccgcgttgctggcgtttttccataggctccgcccccctga<br>cgagcatcacaaaaatcgacgctcaagtcagaggtggcgaaacccgacaggactataaagataccaggcgtttc<br>cccctggaagctccctcgtgcgctctcctgttccgaccctgccgcttaccggatacctgtccgcctttctccct<br>tcgggaagcgtggcgctttctcatagctcacgctgtaggtatctcagttcggtgtaggtcgttcgctccaagct<br>gggctgtgtgcacgaaccccccgttcagcccgaccgctgcgccttatccggtaactatcgtcttgagtccaacc<br>cggtaagacacgacttatcgccactggcagcagccactggtaacaggattagcagagcgaggtatgtaggcggt<br>gctacagagttcttgaagtggtggcctaactacggctacactagaagaacagtatttggtatctgcgctctgct<br>gaagccagttaccttcggaaaaagagttggtagctcttgatccggcaaacaaaccaccgctggtagcggtggtt<br>tttttgtttgcaagcagcagattacgcgcagaaaaaaaggatctcaagaagatcctttgatcttttctacgggg<br>tctgacgctcagtggaacgaaaactcacgttaagggattttggtcatgagattatcaaaaaggatcttcaccta<br>gatccttttaaattaaaaatgaagttttaaatcaatctaaagtatatatgagtaaacttggtctgacagttacc<br>aatgcttaatcagtgaggcacctatctcagcgatctgtctatttcgttcatccatagttgcctgactccccgtc<br>gtgtagataactacgatacgggagggcttaccatctggccccagtgctgcaatgataccgcgagacccacgctc<br>accggctccagatttatcagcaataaaccagccagccggaagggccgagcgcagaagtggtcctgcaactttat<br>ccgcctccatccagtctattaattgttgccgggaagctagagtaagtagttcgccagttaatagtttgcgcaac<br>gttgttgccattgctacaggcatcgtggtgtcacgctcgtcgtttggtatggcttcattcagctccggttccca<br>acgatcaaggcgagttacatgatcccccatgttgtgcaaaaaagcggttagctccttcggtcctccgatcgttg<br>tcagaagtaagttggccgcagtgttatcactcatggttatggcagcactgcataattctcttactgtcatgcca<br>tccgtaagatgcttttctgtgactggtgagtactcaaccaagtcattctgagaatagtgtatgcggcgaccgag<br>ttgctcttgcccggcgtcaatacgggataataccgcgccacatagcagaactttaaaagtgctcatcattggaa<br>aacgttcttcggggcgaaaactctcaaggatcttaccgctgttgagatccagttcgatgtaacccactcgtgca<br>cccaactgatcttcagcatcttttactttcaccagcgtttctgggtgagcaaaaacaggaaggcaaaatgccgc<br>aaaaaagggaataagggcgacacggaaatgttgaatactcatactcttcctttttcaatattattgaagcattt<br>atcagggttattgtctcatgagcggatacatatttgaatgtatttagaaaaataaacaaataggggttccgcgc<br>acatttccccgaaaagtgccacctgac |

#### Supplementary Table 3: Mapping and QC statistics for ChIP-seq datasets used in this study

| Dataset | Library complexity | NSC | RSC | QC | Read Length | Mapped reads | Raw fragments |
|---|---|---|---|---|---|---|---|
| L111-Cas9-sgRNA-N4293-Input | 0.96 | 1.121 | 0.303 | -1 | 2x75 | 84,490,748 | 55,837,397 |
| L120-Cas9-sgRNA-N4293-CTCF | 0.93 | 4.111 | 1.563 | 2 | 2x75 | 34,775,812 | 23,853,267 |
| L121-Cas9-sgRNA-8005-CTCF | 0.9 | 8.302 | 1.909 | 2 | 2x75 | 31,881,148 | 22,583,697 |
| L122-Cas9-sgRNA-12040-CTCF | 0.91 | 8.584 | 1.827 | 2 | 2x75 | 20,212,130 | 14,111,997 |
| L123-Cas9-sgRNA-13004-CTCF | 0.87 | 8.154 | 1.905 | 2 | 2x75 | 27,329,576 | 19,091,732 |
| L124-Cas9-sgRNA-14259-CTCF | 0.91 | 9.061 | 1.932 | 2 | 2x75 | 30,866,786 | 21,744,890 |
| L125-Cas9-sgRNA-14376-CTCF | 0.92 | 9.269 | 1.874 | 2 | 2x75 | 26,398,240 | 19,111,658 |
| L126-Cas9-sgRNA-15776-CTCF | 0.91 | 10.139 | 1.953 | 2 | 2x75 | 29,279,448 | 20,919,316 |

#### Supplementary Table 4: Mapping and QC statistics for RNA-seq datasets used in this study

| Library | Raw fragments | Complexity | Unique | Unique Splices | Multi | Multi Splices | Fraction mapped |
|---|---|---|---|---|---|---|---|
| 12040_1_CTCFvalidation_S1 | 28,644,781 | 0.68 | 20,358,952 | 4,132,800 | 2,424,064 | 640,956 | 0.48 |
| 12040_2_CTCFvalidation_S2 | 28,846,455 | 0.67 | 19,606,123 | 3,975,313 | 2,319,439 | 621,135 | 0.46 |
| 12042_1_CTCFvalidation_S3 | 36,357,512 | 0.66 | 25,770,216 | 5,389,554 | 3,016,467 | 830,405 | 0.48 |
| 12042_2_CTCFvalidation_S4 | 32,696,948 | 0.67 | 22,560,933 | 4,604,545 | 2,604,102 | 697,272 | 0.47 |
| 13004_1_CTCFvalidation_S9 | 28,892,443 | 0.68 | 20,776,068 | 4,282,772 | 2,438,370 | 638,984 | 0.49 |
| 13004_2_CTCFvalidation_S10 | 28,460,588 | 0.68 | 19,421,721 | 3,951,549 | 2,276,571 | 609,239 | 0.46 |
| 13006_1_CTCFvalidation_S11 | 35,962,940 | 0.66 | 25,727,017 | 5,352,693 | 3,000,594 | 821,216 | 0.49 |
| 13006_2_CTCFvalidation_S12 | 31,497,868 | 0.68 | 20,750,328 | 4,296,768 | 2,458,910 | 666,244 | 0.45 |
| 14376_1_CTCFvalidation_S5 | 38,779,897 | 0.66 | 27,758,193 | 5,792,893 | 3,271,621 | 888,843 | 0.49 |
| 14376_2_CTCFvalidation_S6 | 28,806,065 | 0.68 | 18,814,700 | 3,832,676 | 2,155,832 | 583,804 | 0.44 |
| 14377_1_CTCFvalidation_S7 | 30,361,030 | 0.68 | 21,620,017 | 4,310,279 | 2,522,358 | 664,134 | 0.48 |
| 14377_2_CTCFvalidation_S8 | 36,713,802 | 0.66 | 24,774,171 | 5,079,087 | 2,940,671 | 801,139 | 0.46 |
| N4293_1_CTCFvalidation_S13 | 31,819,345 | 0.67 | 22,969,926 | 4,745,722 | 2,665,517 | 724,855 | 0.49 |
| N4293_2_CTCFvalidation_S14 | 31,045,772 | 0.68 | 20,376,549 | 4,265,365 | 2,382,260 | 649,440 | 0.45 |
| N4871_1_CTCFvalidation_S15 | 33,515,599 | 0.65 | 24,289,758 | 5,053,530 | 2,865,219 | 765,455 | 0.49 |
| N4871_2_CTCFvalidation_S16 | 19,600,752 | 0.72 | 13,567,179 | 2,753,987 | 1,553,040 | 416,586 | 0.47 |

**Supplementary Table 5: Mapping and QC statistics for ATAC-seq datasets used in this study**

| Library | Raw fragments | Unique reads | Complexity | chrM reads | chrM fraction | Unique non-chrM reads after dedup | TSS ratio | MACS default peaks | FRiP (MACS) | post IDR (0.05) peaks, ind. Reps | FRIP (IDR) |
|---|---|---|---|---|---|---|---|---|---|---|---|
| Cas9-13004-1 | 52,789,893 | 61,048,866 | 0.82 | 33,514,562 | 0.32 | 50,711,620 | 14.36 | 60,654 | 0.21 | 43,160 | 0.22 |
| Cas9-13004-2 | 18,231,944 | 21,188,935 | 0.87 | 12,106,612 | 0.33 | 18,580,257 | 14.85 | 30,715 | 0.17 | 43,160 | 0.22 |
| Cas9-13006-1 | 28,470,155 | 31,019,550 | 0.87 | 19,482,884 | 0.34 | 27,179,398 | 15.3 | 38,581 | 0.19 | 42,584 | 0.23 |
| Cas9-13006-2 | 21,881,230 | 24,333,878 | 0.86 | 15,785,470 | 0.36 | 21,233,756 | 16.12 | 36,687 | 0.19 | 42,584 | 0.24 |
| Cas9-N4293-1 | 27,588,151 | 35,157,791 | 0.85 | 13,635,924 | 0.25 | 30,100,001 | 12.97 | 47,341 | 0.16 | 43,500 | 0.19 |
| Cas9-N4293-2 | 19,786,147 | 24,311,620 | 0.86 | 11,934,332 | 0.3 | 21,179,734 | 16.86 | 39,213 | 0.21 | 43,500 | 0.25 |
| Cas9-N4371-1 | 30,394,942 | 36,688,985 | 0.84 | 17,928,778 | 0.29 | 31,230,415 | 16.19 | 48,875 | 0.22 | 50,962 | 0.25 |
| Cas9-N4371-2 | 23,086,373 | 27,737,031 | 0.85 | 14,534,148 | 0.31 | 24,025,735 | 17.09 | 45,001 | 0.22 | 50,962 | 0.27 |

### Supplementary Materials & Methods

##### sgRNA targeting CTCF motif library, additional library design methods

In addition to the 4,022 canonical loop anchor CTCF binding sites (Type 0), we added to the screen a small set of 310 sites (Types 1 - 5) that would allow us to test additional hypotheses about the role of CTCF in gene regulation and genome 3D architecture. For each hypothesis, we started with 100 candidate sites, and as before, filtered out the ones with ≥2 sgRNAs passing filtering criteria. However, upon discovering the dominant effect of confounding off-target activity in the CTCF motif screen, which was similarly dominant among sites of Types 1 - 5, we decided not to include these additional types in the analyses and figures for the sake of clarity.

**Type 1: Loop anchors without annotated CTCF binding sites annotated using binding preferences obtained from deep learning models for predicting TF binding:** We hypothesized that we could expand the set of binding loop anchors tested by including CTCF sites that fall below the motif-calling threshold used for annotation before [40] but for which formation of the loop might still be occurring through a CTCF-mediated mechanism. We required remaining unannotated loop anchors to be in a TAD with genes with strong growth effects in gene-level knockout screens. Then, within the loop anchors, we annotated likely CTCF "motifs" by identifying subsequences with high importance for binding in a TF binding prediction deep learning model (described below, **Supplementary Materials & Methods**). Importance scores were derived using DeepLIFT with gradient times input [101]; important subsequences were defined as those that for which the cross-correlation of the DeepLIFT gradient times input scores and the CTCF JASPAR motif [102] was large. After guide filtering, we were left with 80 sites.

**Type 2: Rad21 ChIP-exo peaks in TADs with strongest growth genes:** Here we asked whether the growth effects we might observe at CTCF binding sites are due to disruption of CTCF binding, disruption of RAD21 (or more generally the cohesin complex) function or both. Previous work that used ChIP-exo to map the precise binding of CTCF and RAD21 suggested that these 2 proteins occur in specific spacing and orientations [103], allowing us to test this hypothesis. Thus, to identify RAD21-specific binding sites, we used the existing data from that work as follows. Since Tang et al. did not profile ChIP-exo of RAD21 in K562 cells, we started from the ChIP-exo RAD21 sites measured in GM12878 cells. We kept the RAD21 ChIP-exo sites that overlapped ChIP-seq peaks for RAD21 in K562 cells and required them to be within 100 bp to the right of the CTCF motifs from our screen, consistent with the positioning of RAD21 ChIP-exo sites relative to CTCF ChIP-exo sites in Tang et al. We also prioritized RAD21 binding sites annotated as GSB (spikes on both sides of the binding sites), which are more confident ChIP-exo calls. After filtering, we obtained 72 such sites.

**Type 4: CTCF ChIP-seq peaks outside loop anchors within DNase hypersensitive regions:** In order to compare loop-anchor CTCF sites to non-loop anchor CTCF sites, we selected CTCF sites from the latter category within the top 100 TADs containing growth genes subject to the requirement that they overlap K562 DNase hypersensitive regions from the Roadmap Epigenomics data from the University of Washington [51]. We then defined the precise CTCF binding site based on DeepLIFT scores as described above. The final set after filtering contained 82 sites.

**Type 5: CTCF ChIP-seq peaks outside loop anchors outside DNase hypersensitive sites:** This category was defined as above except that CTCF sites were required to not overlap K562 DNase hypersensitive regions. The final set in this case consisted of 76 sites.

For the analyses used here (**Figure 1**), we filtered the library to remove Types 1-5.

**Deep learning models for TF binding prediction used for CTCF library design:**
To select CTCF sites in the category of loop anchors without annotated CTCF binding sites, we trained a deep convolutional neural network (CNN) to predict whether a sequence is a CTCF binding site or an open chromatin region without CTCF, computed importance scores for nucleotides' importance for the model's predictions, and compared the nucleotides weighted by their importance scores to a PWM for CTCF. The positive set in CNN training consisted of the ±500bp sequences around IDR-reproducible black list-filtered ENCODE K562 CTCF ChIP-seq peak summits (ENCODE accession ID ENCSR000DMA) [60]. For the negative set, the ±500bp sequences around Epigenomic Roadmap [51] K562 DNase peak summits that did not overlap any K562 CTCF peaks (including non-reproducible peaks) were used. The training set consisted of chromosomes 3-7, 10-22 and X; the validation set (used for hyper-parameter tuning) consisted of chromosomes 8 and 9, and the test set consisted of chromosomes 1 and 2. Sequences were one-hot encoded as 4*1000 binary matrices following previously established practices [104,105]. "N" bases were encoded as zeros. We separately encoded each sequence and its reverse complement.

We used an architecture featuring three convolutional layers, with each followed by a rectified linear unit (ReLU), followed by a max-pooling layer. The convolutional filters of the first layer can be interpreted as picking up sequence patterns revealing whether a peak is a CTCF peak or a DNase peak without CTCF, the filters in the following layers identify combinations of those patterns, and the max-pooling layer encodes the assumption that a single sequence pattern combination should not occur multiple times within a short region. The first convolutional layer had sixty (4*15) filters with stride 1*1, the second convolutional layer had 60 (1*15) filters with stride 1*1, and the third convolutional layer had 15 (1*15) filters with stride 1*1. Each layer used a dropout rate of 0.2. The max-pooling layer was of size 1*35 and stride 1*35. The max-pooling layer was followed by a sigmoid. The model was trained using Keras version 0.3.2 [106] with stochastic gradient descent with Nesterov momentum 0.85, learning rate 0.01, and batch size 200. The model was trained for 47 epochs. Weights were initialized from a pre-trained model with the same hyper-parameters and the negative set randomly down-sampled to be the size of the positive set, where the model was trained for 100 epochs. Weights for pre-training were initialized using Keras's He normal initializer [106,107].

To identify regions within ChIP-seq peaks that are important for making positive predictions, we scored the importance of every nucleotide in each positive example using DeepLIFT with gradient times input, which computes the product of each input and gradient with respect to that input (Shrikumar, Greenside, & Kundaje, 2017). Since most CTCF ChIP-seq peaks that were correctly predicted had at least one region with high DeepLIFT scores and we wanted to select less than one hundred guides, we filtered the regions with high DeepLIFT scores by cross-correlating the scores starting at each index within the sequence with the log-odds of the CTCF PWM from JASPAR [102], where we used a pseudo-count of 0.0001 and a background of 52% GC content when computing the log-odds. This was procedure was carried out for each sequence and its reverse complement, and the top two motif hits across both were retained. (Note that some of these motif hits would not be identified by scanning the sequence for the CTCF motif because the regions with important DeepLIFT scores are not always those with the best matches to the CTCF motif.) Motif hits within 20 bp of a higher-scoring motif hit as well as those with log-odds scores ≤0.5 were removed. Motif hits in peaks without a previously identified CTCF motif hit were retained; the intuition is that these sequences are imperfect matches to the CTCF motif that are missed by PWM scanning.
